# Elevating levels of neuronal MCU in the hippocampus enhances mitochondrial calcium uptake and respiratory efficiency proportional to demand

**DOI:** 10.64898/2026.04.13.718264

**Authors:** Mikel L. Cawley, Ryan N. Montalvo, Mason L. Wheeler, Lucy L. Turner, Jessica Pfleger, Zhen Yan, Shannon Farris

**Author notes:** Department of Integrative Physiology, Baylor College of Medicine, Houston, Texas (Jessica Pfleger).

## Abstract

Mitochondrial calcium signaling integrates energy needs with energy production, amplifying or suppressing mitochondrial respiration in response to activity demand. Neuronal activity is tightly ATPcoupled to increases in mitochondrial calcium uptake, which stimulate the tricarboxylic acid cycle (TCA) and activate calcium-dependent enzymes important for ATP production via oxidative phosphorylation. The mitochondrial calcium uniporter (MCU) is the predominant source of matrix calcium and is differentially expressed across neuronal cell types, suggesting cell-type-specific differences in the coupling of activity-driven calcium levels and mitochondrial respiration. Here, we investigated whether elevating MCU expression enhances mitochondrial calcium uptake and oxidative phosphorylation in the hippocampus. We report that hippocampal mitochondria overexpressing MCU take up calcium at a faster rate without increased sensitivity to calcium overload. By modeling in vivo supply and demand, we found that hippocampal mitochondria overexpressing MCU are more efficient than control mitochondria at responding to increased bioenergetic demand. These findings reveal a role for MCU in modulating mitochondrial calcium uptake and boosting mitochondrial respiration under increasing demand, which contributes to our understanding of how specific cell types may adapt to different bioenergetic demands.

## 1 INTRODUCTION

Neurons are particularly sensitive to mitochondrial dysfunction because of their high energy demands and complex cytoarchitecture that requires mitochondria to be properly distributed and maintained over long distances. Consequently, mitochondrial dysfunction is implicated in several neurological and neurodegenerative disorders. It is well established that certain brain regions are more susceptible to neuropathology, and one theory is that their differential energetic burden makes them more vulnerable. Therefore, investigating how mitochondria regulate energy production to support diverse neuronal metabolic demands is critical for understanding regional vulnerability to mitochondrial dysfunction (Mattson, 2007; Calì, Ottolini and Brini, 2012; Verma, Lizama and Chu, 2022).

The mitochondrial calcium uniporter (MCU) complex regulates energy production in neurons, linking cytosolic and mitochondrial calcium levels to mitochondrial respiration (Nichols, Pavlov and Robertson, 2018; Groten and MacVicar, 2022; Stoler *et al*., 2022; Li *et al*., 2024). The MCU complex is the predominant source of calcium entry into the mitochondrial matrix, which activates calcium-dependent dehydrogenases (isocitrate, oxoglutarate, and pyruvate dehydrogenases) and the ETC complex V, thereby stimulating the proton motive force and driving ATP synthesis (Kamer and Mootha, 2015; De Mario *et al*., 2023; Cartes-Saavedra, Ghosh and Hajnóczky, 2025). The main pore-forming subunit, MCU, is located on the inner mitochondrial membrane (IMM) and gated by the calcium-sensitive EF-hand regulators mitochondrial calcium uptake 1(MICU1), MICU2, and MICU3 (Perocchi *et al*., 2010; De Stefani *et al*., 2011; Patron *et al*., 2014; Paillard *et al*., 2017). The ratio of MICU1 to MCU varies across tissue types (Paillard *et al*., 2017; Farris *et al*., 2019) and differentially regulates pore conductivity (Paillard *et al*., 2017). We previously reported that the hippocampal subregion CA2 shows enrichment of several components of the MCU complex relative to neighboring subregions, including the MCU pore, MICU1, and MCUR1 (Farris *et al*., 2019). In particular, MCU is enriched in CA2 distal dendrites (Pannoni *et al*., 2023), where it regulates synaptic plasticity (Pannoni *et al*., 2025) and results in higher basal mitochondrial matrix calcium levels than in CA2 proximal dendrites (Alsalman *et al*., 2025). Collectively, these data suggest that functionally distinct hippocampal circuits harbor unique mitochondrial populations with different capacities for calcium handling and energy production. Neurons expressing higher endogenous MCU levels showed relatively greater mitochondrial calcium uptake than neurons with lower MCU levels (Fecher *et al*., 2019; Groten and MacVicar, 2022). Preventing MCU-mediated mitochondrial calcium uptake either pharmacologically in live brain slices (Stoler *et al*., 2022) or via shRNA knockdown in culture (Ashrafi *et al*., 2020; Ghosh *et al*., 2025) uncoupled activity-driven increases in NADH or ATP production, respectively. Together, these findings support a role for varied neuronal MCU expression in tuning cell-type or circuit-specific bioenergetic profiles.

MCU-dependent calcium uptake could exert feed-forward regulation of oxidative phosphorylation by increasing matrix calcium, dehydrogenase activity, and TCA cycle flux. However, calcium uptake must be balanced by calcium efflux, or it risks activating the mPTP and triggering cell death pathways. Therefore, the tight coupling between mitochondrial calcium uptake and mitochondrial function was predicted to be essential for neuronal metabolism. However, the global loss of MCU in nonsynaptic mitochondria from rat brain did not result in any deficits in oxidative phosphorylation compared with wild type (Hamilton *et al*., 2018). In contrast, directly driving oxidative phosphorylation via MCU-dependent calcium uptake in brain mitochondria was stimulatory at low concentrations and inhibitory at high concentrations (Pandya, Nukala and Sullivan, 2013). The dose-dependent inhibitory effect of calcium on oxygen consumption was not attributed to mPTP opening, but an acute response to supraphysiological calcium with long-term consequences on complex I function (Pandya, Nukala and Sullivan, 2013). Therefore, it remains unclear whether differential MCU expression scales mitochondrial calcium uptake with respiratory capacity.

Here, we aimed to determine the bioenergetic consequences of enriched MCU expression in the hippocampus. We used adeno-associated viral vectors (AAVs) to overexpress the MCU pore, or GFP as the control, in hippocampal neurons and isolated mitochondria to assess mitochondrial calcium kinetics and respiration. Our findings indicate that elevated MCU pore expression is sufficient to increase the rate of mitochondrial calcium uptake without increasing sensitivity to calcium overload. Furthermore, elevated MCU expression not only enhanced mitochondrial respiration across varying substrate conditions but also increased the efficiency of ATP regeneration in response to growing energetic demand. These findings strengthen evidence for the differential MCU expression as a mechanism to scale mitochondrial calcium and respiration in proportion to cell-type-specific bioenergetic demands. Differential expression of MCU complex proteins may reflect a general mechanism by which specific cell types adapt mitochondrial calcium handling and energetic load, which may influence vulnerability to mitochondrial dysfunction in neurodegenerative disease (Garcia-Casas *et al*., 2023; Twyning *et al*., 2024).

## 2 RESULTS

### 2.1 Neuronal mitochondria overexpressing MCU take up calcium at a higher rate

To determine if mitochondrial calcium dynamics are enhanced in neuronal mitochondria with elevated MCU levels, we bilaterally infused adeno-associated viral vectors (AAV) in the hippocampus of adult male and female mice to overexpress MCU (AAV9-hSyn-mMCU-2A-eGFP-WPRE), or GFP (AAV9-hSyn-eGFP-WPRE) as a control (**Fig. 1A**). Two to three weeks after infusion, MCU expression was increased by ∼70% in mitochondrial fractions from hippocampi of MCU overexpression (MCU OE) compared to GFP controls (**Fig. 1B,C**, two-tailed unpaired t–test, t(34) = 6.4, p < 0.0001, N = 18 mice/group). There was a small but not significant decrease in the expression of MICU1 in MCU OE mitochondrial fractions compared to GFP controls (**Fig. 1B,D**, avg. norm. MICU1 fold change (F.C.) to GFP ±SEM, GFP: 1.00 ±0.06; MCU OE: 0.87 ±0.05; two-tailed unpaired t–test, p > 0.05, N = 8 mice/group). There was a significant decrease in the ratio of MICU: MCU protein expression in MCU OE fractions compared to GFP controls (**Fig. 1E**, two-tailed unpaired t–test, p < 0.002, N = 8 mice/group). MCU overexpression was also validated by immunohistochemistry and resulted in a ∼5-fold increase in MCU fluorescence intensity in CA1 neurons compared to GFP-expressing control (**S.1,** two-tailed unpaired t-test; p ≤ 0.0001, N = 6 mice/group).

**Figure 1:**
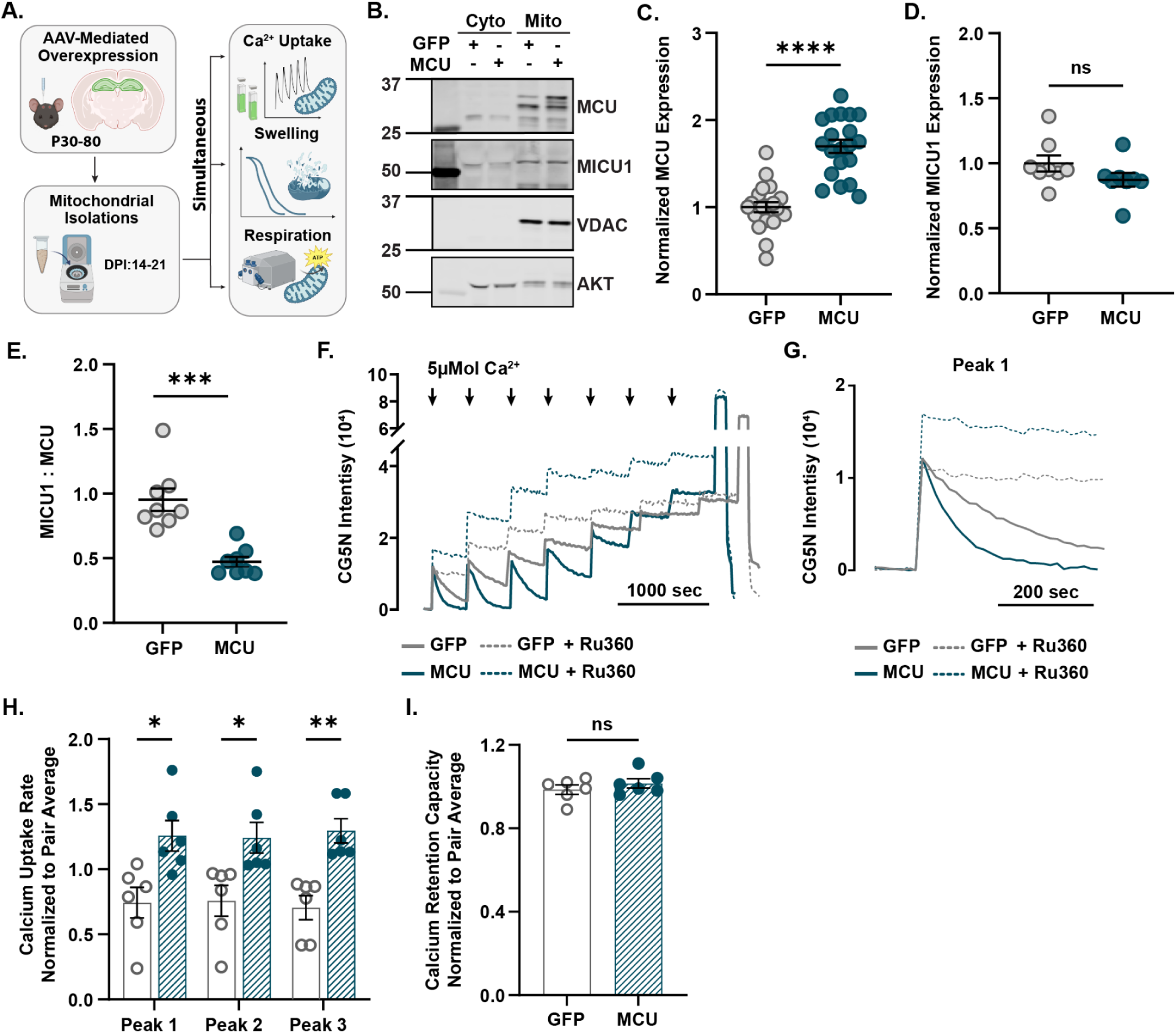
MCU overexpression in hippocampal neurons increases mitochondrial calcium uptake without altering calcium retention capacity relative to GFP-expressing controls. A) MCU or GFP was bilaterally overexpressed in hippocampal neurons of adult male and female mice. Hippocampi were homogenized from MCU- and GFP-overexpressing pairs 14-21 days post-infusion (dpi). Standard differential centrifugation was used to isolate crude mitochondrial fractions for calcium and bioenergetic assays. Mitochondrial isolations were performed on MCU OE and GFP pairs to enable tandem bioenergetic and calcium assays. B) Representative immunoblot using antibodies against MCU, MICU1, VDAC, and AKT in mitochondria isolated from hippocampi overexpressing GFP or MCU. VDAC was used as a mitochondrial loading control. AKT was used as a cytosolic loading control. Full uncropped membranes are shown in the supplementary materials (**S.11**). C) Quantification shows MCU protein expression normalized to mitochondrial content (VDAC). Densitometry is reported as a fold change (F.C) relative to GFP controls (avg. F.C MCU to GFP ±SEM, GFP: 1.00 ±0.07; MCU OE: 1.70 ±0.08; two-tailed unpaired t–test, t(34) = 6.4, ****p < 0.0001, N = 18 mice/group). The integrated density of each MCU band was summed to reflect differences in the amount of endogenous and exogenous MCU expression between groups, as MCU was previously shown to oligomerize and form large interactomes with the IMM due to higher levels of MCU activity (Bharat *et al*., 2023; Delgado de la Herran *et al*., 2024). Full uncropped membranes corresponding to quantification are shown in the supplementary materials (**S.12,13**). D) Quantification shows MICU1 protein expression normalized to mitochondrial content (VDAC). Densitometry is reported as a fold-change relative to GFP controls (avg. F.C MICU1 to GFP ±SEM, GFP: 1.00 ±0.06; MCU OE: 0.87 ±0.05; two-tailed unpaired t–test, p = 0.15, N = 8 mice/group). Full uncropped membranes corresponding to quantification are shown in the supplementary materials (**S.13,14**). E) Quantification shows the ratio of MICU1 to MCU protein expression. MICU1:MCU is reported as the ratio of density for MICU1 and MCU (avg. MICU1:MCU ±SEM, GFP: 0.95 ±0.09; MCU OE: 0.47 ±0.04; two-tailed unpaired t–test, ***p = 0.0002, N = 8 mice/group). Each point represents an independent biological replicate. F) Representative calcium uptake trace from isolated hippocampal mitochondria (0.125mg/mL of protein) expressing GFP (grey) or overexpressing MCU (teal) in the absence (solid) and presence (dashed) of MCU blocker Ru360 (20μM). Mitochondrial calcium uptake rate was measured by the decrease in extramitochondrial calcium via the fluorescent indicator Calcium Green-5N (CG5N). Sequential 6 nmol Ca^2+^ pulses were added every 7 minutes (black arrows) until CG5N intensity plateaued. The final peak shows fluorescence from a saturating Ca2+ pulse (1 M), followed by EGTA (0.5 M), to determine the dynamic range. The calcium retention capacity was calculated as the cumulative Ca^2+^ load taken up prior to the plateau. Trace represents a single GFP and MCU OE paired run conducted on the same day. CG5N intensity values for each trace are background-subtracted and normalized to baseline. G) Representative Calcium Green-5N trace showing the slope of single 6 nmol Ca^2+^ pulses used to extrapolate calcium uptake rates from mitochondria overexpressing GFP (grey) and MCU (teal) in the absence (solid) and presence (dashed) of Ru360 (20μM). H) Quantification of mitochondrial calcium uptake rates for three sequential Ca2+ pulses (Final: 30 μmol Ca^2+^) from isolated mitochondria expressing GFP or overexpressing MCU. Calcium uptake (Ca²⁺nmol·s⁻¹·mg⁻¹) rates were derived from the slope of extramitochondrial calcium decline, measured by loss in Calcium Green-5N fluorescence intensity per peak. Calcium uptake rates are normalized to the average rate for each pair (avg. peak 1 ±SEM: GFP: 0.7 ±0.1; MCU OE: 1.3 ±0.1; avg. peak 2 ±SEM: GFP: 0.8 ±0.1; MCU OE:1.2 ±0.1; avg. peak 3 ±SEM: GFP: 0.7 ±0.09; MCU OE: 1.3 ±0.09; multiple unpaired t-tests with Holm-Ṧídák’s correction, *p ≤ 0.05, **p ≤ 0.01, N = mice/group). I) Quantification shows calcium retention capacity from isolated mitochondria expressing GFP or overexpressing MCU. Calcium retention capacity (Ca²⁺nmol·mg⁻¹) represents the total amount of Ca^2+^ taken up before Calcium Green-5N fluorescence intensity plateaus. Calcium retention capacity was normalized by the average capacity of each pair (avg. CRC ±SEM: GFP: 0.98 ±0.22; MCU OE: 1.02 ±0.22; two-tailed unpaired t–test, t(10) = 0.96, p > 0.05, N = 6mice/group).

We then quantified the rate and capacity of mitochondrial calcium uptake in isolated mitochondrial fractions from mice overexpressing MCU or GFP. High-resolution spectrometry and the calcium-sensitive indicator Calcium Green-5N (CG5N) were used to measure the uptake rate and calcium retention capacity (CRC) of neuronal mitochondria exposed to increasing extramitochondrial calcium concentrations over time (**Fig. 1F**). The MCU inhibitor Ru360 (de J García-Rivas *et al*., 2006) was used to confirm that the source of calcium uptake was through the MCU pore (**Fig. 1F,G**). To account for technical variation between day-to-day runs, calcium uptake rates and capacity were normalized to the average for each MCU OE and GFP-expressing pair. Raw traces for each pair are available in supplemental materials (**S.2)**. Raw calcium uptake rates per peak (Ca²⁺nmol·s⁻¹·mg⁻¹) for each pair are presented in supplemental materials (**S.3**). There was a significant increase in mitochondrial calcium uptake rate per peak for MCU OE mitochondria compared to GFP-expressing controls (**Fig. 1H**, multiple unpaired t-tests with Holm-Ṧídák’s correction, p ≤ 0.05, N = 6 mice/group). There was no significant effect of MCU OE on calcium retention capacity compared with GFP-expressing controls, suggesting that MCU OE mitochondria are more efficient calcium buffers than GFP-expressing controls (**Fig. 1I**, avg. CRC ±SEM: GFP: 0.98 ±0.22; MCU OE:1.02 ±0.22, two-tailed unpaired t-test; p > 0.05, N = 6 mice/group). Raw values for calcium uptake capacity (Ca²⁺nmol·mg⁻¹) for each pair are presented in supplemental materials (**S.3**). These results demonstrate that elevating MCU levels enhances the rate of mitochondrial calcium uptake, but the capacity to retain calcium remains unchanged compared to GFP-expressing controls.

### 2.2 Overexpressing MCU in neurons does not increase sensitivity to calcium overload

A high influx of calcium into the mitochondrial matrix triggers opening of the mitochondrial permeability transition pore (mPTP), increasing IMM permeability and eventually rupturing the outer mitochondrial membrane (Bernardi *et al*., 2023). While we did not detect a difference in capacity, overexpressing MCU by ∼70% could increase vulnerability to mPTP opening by elevating matrix calcium levels. To determine whether neuronal mitochondria overexpressing MCU are more susceptible to mitochondrial permeability transition. Calcium overload was inferred from increasing CG5N fluorescence intensity over time as exposure to high levels of calcium led to swelling and release of mitochondrial calcium (**Fig. 2A**). To capture mitochondrial permeability transition and potential differences in sensitivity to calcium overload, MCU OE- and GFP-isolated mitochondria were challenged with high concentrations of calcium and the relative difference in CG5N intensity (ΔF/F_0_) over time was compared (**Fig. 2B**). Ru360 was used to confirm that the observed loss of CG5N intensity was due to MCU-dependent mitochondrial calcium uptake (**Fig. 2CD**). Cyclosporin A was used to inhibit mPTP and confirm that increased extramitochondrial calcium levels depended on mPTP activation (**S.4**). There was no significant main effect of the group on CG5N Peak ΔF/F_0_ in the presence of calcium (**Fig. 2E**, mixed-effects two-way ANOVA; F(1, 18) = 0.3014, p = 0.5898, N= 4-7 mice/group) or interaction between group x treatment (F(1, 18) = 0.02242, p = 0.8827). There was a significant main effect of treatment on GFP and MCU OE CG5N Peak ΔF/F_0_ in the presence of Ru360, which reduced calcium overload for both groups (**Fig. 2E**, mixed-effects two-way ANOVA; F(1, 18) = 62.18, p < 0.0001; Fisher’s LSD post hoc tests are shown on the plot). These results confirm that increasing extramitochondrial calcium levels depended on MCU-mediated mitochondrial calcium uptake and mPTP activation, but MCU overexpression did not affect sensitivity to calcium overload compared with GFP-expressing controls.

**Figure 2:**
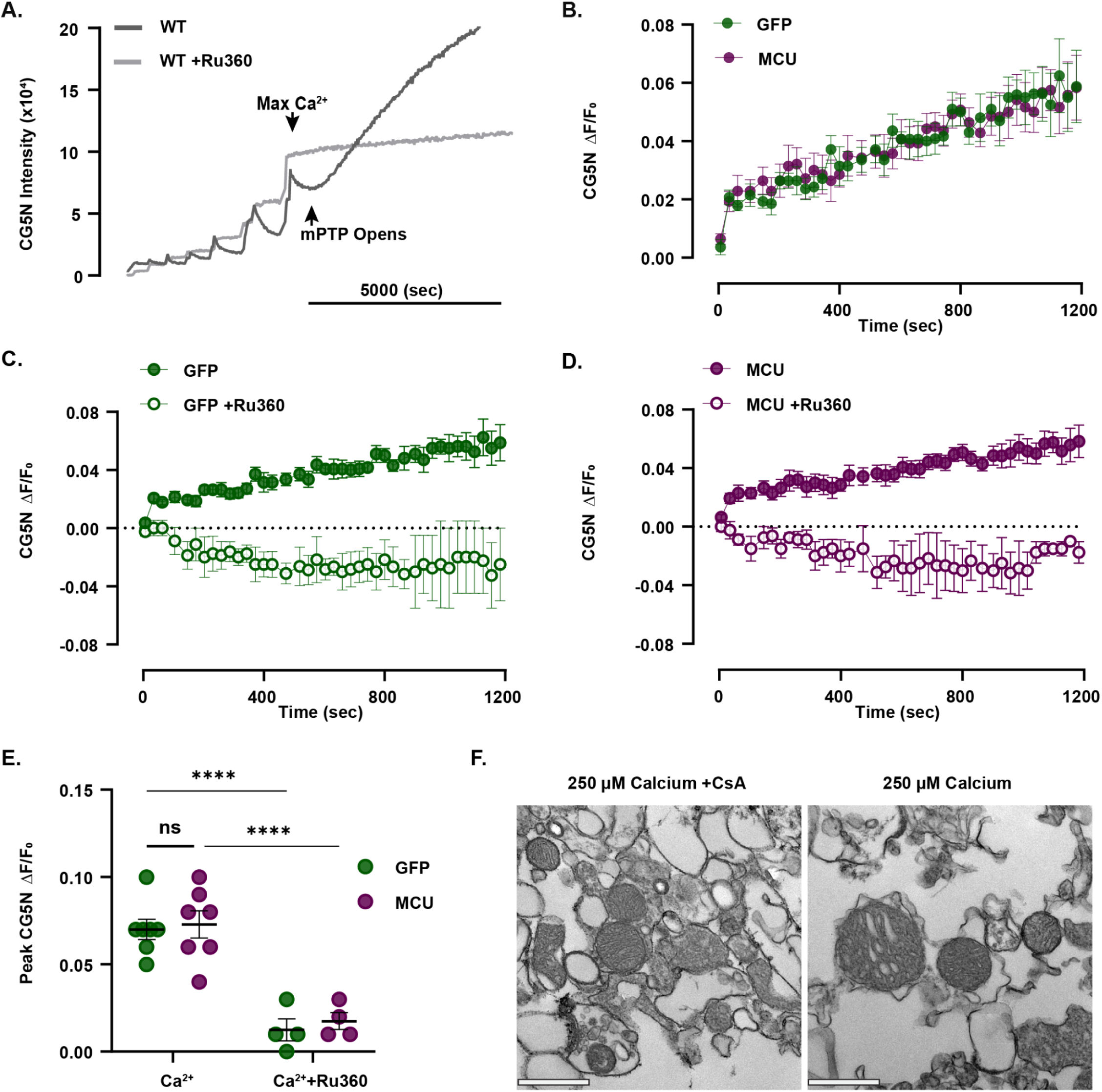
MCU overexpression in hippocampal neurons does not increase mPTP sensitivity to calcium overload compared to GFP-expressing controls. A. Representative trace of Calcium Green-5N (CG5N) intensity during CRC assay of isolated mitochondria (0.125μg/μL) from the hippocampus of a WT mouse in the absence (dark grey) and presence (light grey) of MCU blocker Ru360 (20μM). Sequential Ca^2+^ pulses were added until Ca^2+^ could no longer be taken up (Max Ca^2+^: 60µM), which induced calcium overload (mPTP opening) and resulted in a substantial increase in CG5N intensity in the absence of Ru360. CG5N intensity values for each trace are background-subtracted and normalized to baseline. B. Mean CG5N ΔF/F_0_ trace from MCU OE and GFP-expressing mitochondria. Mean CG5N ΔF/F_0_ traces reflect the average trace per group recorded over 1200 seconds from several independent assays in response to large calcium (range: 35-300μM) pulses (avg. ΔF/F_0_ ±SEM: GFP: 0.39 ±0.14; MCU OE: 0.39 ±0.13; N = 7 mice/group). C. Mean CG5N ΔF/F_0_ trace from GFP-expressing mitochondria in the absence or presence of Ru360 (20μM). Mean CG5N ΔF/F_0_ traces reflect the average trace per treatment recorded over 1200 seconds (avg. ΔF/F_0_ ±SEM: GFP: 0.39 ±0.14; N = 7, GFP+Ru360: -0.02 ±0.01; N =4). D. Mean CG5N ΔF/F_0_ trace from MCU OE mitochondria in the absence or presence of Ru360. Mean CG5N ΔF/F_0_ traces reflect the average trace per treatment recorded over 1200 seconds (avg. ΔF/F_0_ ±SEM: MCU OE: 0.39 ±0.13; N = 7, MCU OE +Ru360: -0.02 ±0.02; N = 4). E. Quantification for peak CG5N ΔF/F_0_ values from MCU OE and GFP-expressing mitochondria in the absence or presence of Ru360. There was no main effect of group on peak CG5N ΔF/F_0_ in the presence of calcium (mixed-effects model two-way ANOVA; F(1, 18) = 0.3014, p = 0.5898) or interaction between group x calcium (mixed-effects model two-way ANOVA; F(1, 18) = 0.02242, p = 0.8827). There was a significant main effect of Ru360 treatment on CG5N peak ΔF/F_0_ in the presence of calcium (mixed-effects two-way ANOVA; F(1, 18) = 62.18, ****p < 0.0001; Fisher’s LSD). N = 4-7 mice/group). F. Representative TEM images of MCU OE isolated mitochondria recovered from mitochondrial suspensions post calcium overload in the absence or presence of mPTP inhibitor Cyclosporin A. Quantification for peak CG5N ΔF/F_0_ and Mean CG5N ΔF/F_0_ trace in the absence or presence of Cyclosporin A are found in supplemental materials (**S.4**). Scale = 500 nm.

### 2.3 Neuronal mitochondria with elevated MCU levels increase efficiency proportional to energetic demand

Matrix calcium levels can stimulate ATP synthase or other enzymatic reactions that increase electron availability by producing NADH and FADH_2_ (Bochkova *et al*., 2025). To determine if overexpressing MCU enhances mitochondrial respiration in response to demand, we used high resolution respirometry (Oroboros, Innsbruck, Austria) to quantify oxygen consumption rates (*J*O_2_). To determine whether any effects of elevated MCU are responsive to fluctuating energetic demand (Hamilton *et al*., 2018; Ashrafi *et al*., 2020; Stoler *et al*., 2022; Ghosh *et al*., 2025), we used the creatine kinase energetic clamp to assess oxygen flux under a physiologically relevant ATP:ADP ratio (Fisher-Wellman *et al*., 2018). The creatine kinase energetic clamp uses the phosphocreatine/creatine kinase reaction to control ATP:ADP ratios, effectively “clamping” the energetic state. To model *in vivo* supply and demand, we compared respiratory conductance (the slope of *JO*_2_) and ATP free energy (ΔG_ATP_). Higher conductance indicates lower resistance in the electron transport chain, enabling more efficient electron flow and increased ATP production in response to rising energy demand.

To differentiate the contributions of complex I and complex II, substrates malate, pyruvate, and glutamate (MPG, complex I) or succinate, and the complex I inhibitor rotenone (SR, complex II), were used independently to stimulate respiration. To account for potential technical variation between day-to-day runs, the raw *J*O_2_ values for each MCU OE and GFP-expressing pair were normalized to the respective pair average. Raw *J*O_2_ values for each pair are provided in supplemental materials (**S.5,6**). MCU OE mitochondria stimulated by MPG had significantly higher maximal (ΔG_ATP_ -12.94) oxygen consumption rates than GFP mitochondria (**Fig. 3A**, multiple unpaired t-tests with Holm-Šídák’s correction, p ≤ 0.05, N = 10 mice/group). The slope of respiratory conductance (efficiency in response to demand) was also significantly higher in MCU OE mitochondria stimulated by MPG than in GFP mitochondria (**Fig. 3A**, multiple unpaired t-tests with Holm-Šídák’s correction; p ≤ 0.05; N = 10 mice/group). The linear relationship between *J*O_2_ and ΔG_ATP_, measured by phosphocreatine titration (**Fig. 3B**), suggests that MCU OE enhances the efficiency of complex I-linked mitochondrial respiration in response to increased energetic demands. MCU OE mitochondria stimulated with MPG substrates showed a significant increase in PCr titrated *J*O_2_ compared to GFP mitochondria (**Fig. 3C**; multiple unpaired t-tests with Holm-Šídák’s correction, p ≤ 0.05, N = 10 mice/group).

**Figure 3.**
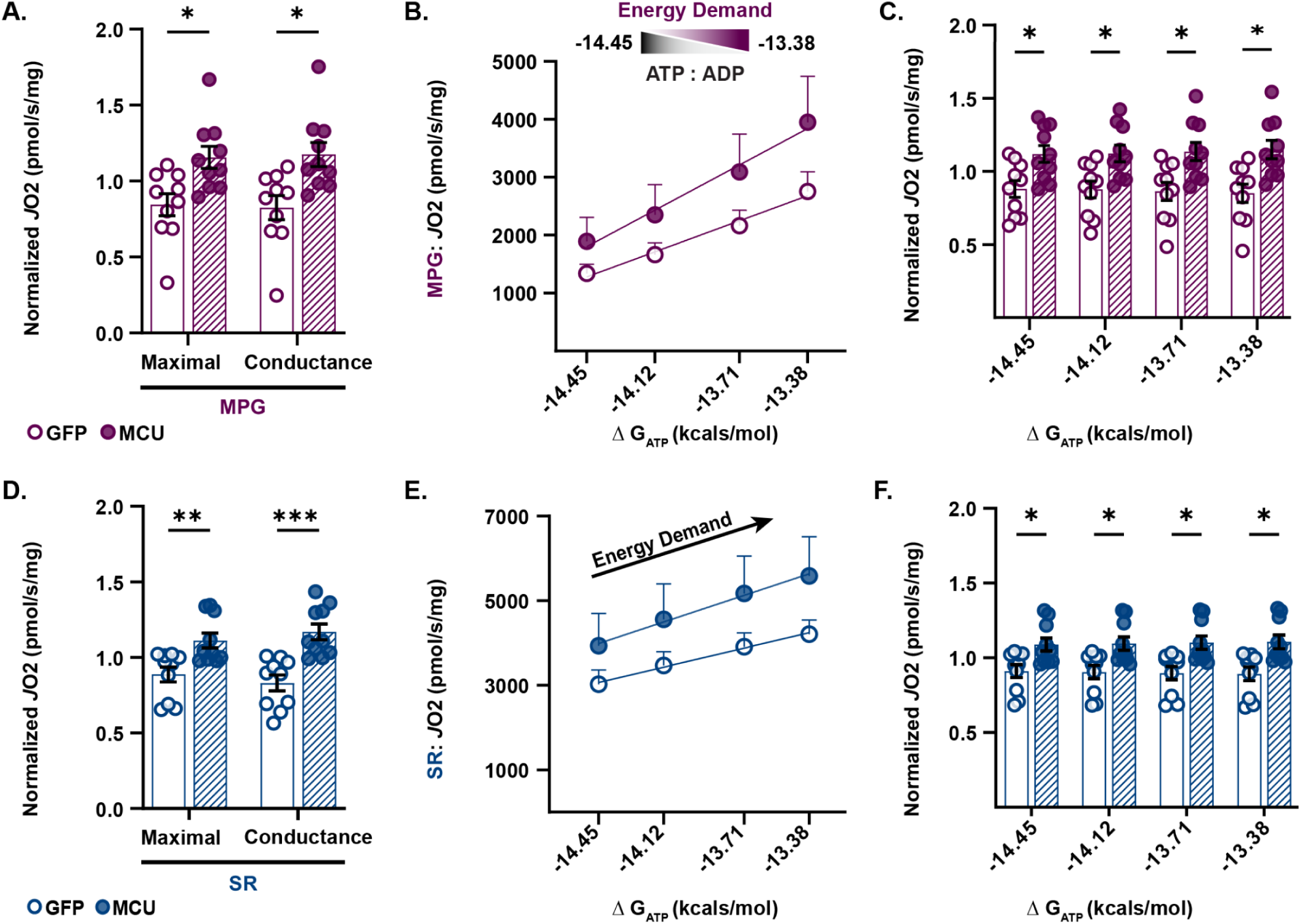
MCU overexpression enhances electron transport chain efficiency under greater energetic demand than in GFP-expressing controls. _A)_ Complex I-linked normalized *J*O_2_ (pmol O₂·s⁻¹·mg⁻¹ mitochondrial protein) energized by malate, pyruvate, and glutamate for maximal and respiratory conductances measured in isolated mitochondria (50µg) expressing GFP or overexpressing MCU. Oxygen consumption rates and respiratory conductances are normalized by the average rate of each pair (norm. max *J*O_2_ ±SEM, GFP: 0.84 ±0.07; MCU OE: 1.16 ±0.07; norm. cond ±SEM, GFP: 0.82 ±0.08; MCU OE: 1.18 ±0.08; multiple unpaired t-tests; all comparisons *p ≤ 0.05, corrected Holm-Ṧídák’s tests, N = 10 mice/group). Oxygen consumption rates are normalized by the average rate of each pair. Raw values are presented in supplemental materials (**S.5,6**). B) The linear relationship between complex I-linked *J*O_2_ (pmol O₂·s⁻¹·mg⁻¹) and ΔG_ATP_ during PCr titration in isolated mitochondria expressing GFP or overexpressing MCU stimulated with MPG. The slope of *J*O_2_ vs. ΔG_ATP_ was quantified and reported as conductance. Each point reflects the mean ±SEM. N = 10 mice/group. _C)_ Complex I-linked normalized *J*O_2_ (pmol O₂·s⁻¹·mg⁻¹ mitochondrial protein) under increasing ΔG_ATP_ in isolated mitochondria expressing GFP or overexpressing MCU stimulated with MPG (avg. norm. *J*O_2_ -14.45 ΔG_ATP_ ±SEM, GFP: 0.88 ±0.06; MCU OE: 1.12 ±0.06; avg. norm. *J*O_2_ -14.12 ΔG_ATP_ ±SEM, GFP: 0.88 ±0.06; MCU OE: 1.12 ±0.06; avg. norm. *J*O_2_ -13.71 ΔG_ATP_ ±SEM, GFP: 0.86 ±0.06; MCU OE: 01.14 ±0.06; avg. norm. *J*O_2_ -13.38 ΔG_ATP_ ±SEM, GFP: 0.85 ±0.06; MCU OE: 1.15 ±0.06; multiple unpaired t-tests; all comparisons *p ≤ 0.05, corrected Holm-Ṧídák’s tests, N = 10 mice/group). Raw values are presented in supplemental materials (**S.6**). D) Complex II-linked normalized *J*O_2_ (pmol O₂·s⁻¹·mg⁻¹ mitochondrial protein) energized by succinate and the complex I inhibitor rotenone for maximal and respiratory conductances measured in isolated mitochondria (50µg) expressing GFP or overexpressing MCU. Each point represents an independent biological replicate from a GFP and MCU OE pair (avg. norm. max *J*O_2_ ±SEM, GFP: 0.89 ±0.05; MCU OE: 1.11 ±0.05; avg. norm. cond ±SEM: GFP: 0.83 ±0.05; MCU OE: 1.17 ±0.05; multiple unpaired t-tests; all comparisons **p ≤ 0.01,***p ≤ 0.001, corrected Holm-Ṧídák’s tests, N = 10 mice/group). Oxygen consumption rates and respiratory conductances are normalized by the average rate of each pair. Raw values are presented in supplemental materials (**S.5,6**) E) The linear relationship between complex II-linked *J*O_2_ (pmol O₂·s⁻¹·mg⁻¹) and ΔG_ATP_ during PCr titration in isolated mitochondria expressing GFP or overexpressing MCU stimulated with SR. The slope of *J*O_2_ vs. ΔG_ATP_ was quantified and reported as conductance. Each point reflects the mean ±SEM. N = 10 mice/group. _F)_ Complex II-linked normalized *J*O_2_ (pmol O₂·s⁻¹·mg⁻¹ mitochondrial protein) under increasing ΔG_ATP_ in isolated mitochondria expressing GFP or overexpressing MCU stimulated with SR (avg. norm. *J*O_2_ -14.12 ΔG_ATP_ ±SEM, GFP: 0.90 ±0.04; MCU OE: 1.10 ±0.04; avg. norm. *J*O_2_ -13.71 ΔG_ATP_ ±SEM, GFP: 0.90 ±0.04; MCU OE: 1.10 ±0.04; avg. norm. *J*O_2_ -13.38 ΔG_ATP_ ±SEM, GFP: 0.89 ±0.05; MCU OE: 1.11 ±0.05; multiple unpaired t-tests; all comparisons *p ≤ 0.05, corrected Holm-Ṧídák’s tests, N = 10 mice/group). Raw values are presented in supplemental materials (**S.6**).

Similarly, compared to GFP mitochondria, MCU OE mitochondria showed increased maximal oxygen consumption rates and respiratory conductance slopes during complex II-linked respiration when stimulated by SR (**Fig. 3D**, multiple unpaired t-tests with Holm-Šídák’s correction, p ≤ 0.001, N = 10 mice/group). The slope of complex II-linked *J*O_2_ and ΔG_ATP_ in response to greater energetic demands was also increased in MCU OE mitochondria relative to GFP mitochondria (**Fig. 3E**). MCU OE mitochondria stimulated with SR substrates showed significant increases in PCr titrated *J*O_2_ compared to GFP mitochondria (**Fig. 3F**, multiple unpaired t-tests with Holm-Ṧídák’s correction, p ≤ 0.05, N = 10 mice/group)

Notably, we found a significant difference between MCU OE and GFP normalized state I (mitochondria only) I *J*O_2_ values, suggesting mitochondria in the presence of buffer exhibit differences in endogenous substrate levels between groups (**S7.A,B;** two-tailed unpaired t-test; p ≤ 0.05, p ≤ 0.01; N = 10 mice per group). Furthermore, we found a significant increase in the complex II-linked non-phosphorylating respiratory state in MCU OE compared to GFP, due to the combination of an oxidizable substrate, succinate, and the complex I inhibitor rotenone (**S7.C,D,** two-tailed unpaired t-test; p ≤ 0.01, N = 10 mice/group). Collectively, these findings reveal that MCU overexpression significantly improves the efficiency of oxidative phosphorylation in response to bioenergetic demand compared to isolated mitochondria expressing GFP.

### 2.4 MCU enrichment does not increase expression levels of the representative ETC complex proteins

Elevated MCU pore expression may promote remodeling of ETC complexes, thereby enhancing oxidative phosphorylation activity and respiratory efficiency. The activity of individual ETC complexes is influenced by the expression levels of complex proteins, post-translational modifications, and/or rearrangements that form supercomplexes, which fine-tune oxidative phosphorylation (Bochkova *et al*., 2025). To determine whether overexpressing MCU increased the expression levels of a subset of electron transport chain complex proteins, we used Western blotting to quantify the relative levels of ETC complex subunits in mitochondrial fractions from a subset of isolations that underwent tandem respiratory assays (**Fig. 4A**). We found no significant difference in the levels of ETC complex I subunit NDUFB8, complex II subunit SDHB, complex III subunit UQCR2, complex IV subunit MTCO1, and complex V subunit ATP5A in mitochondrial fractions overexpressing MCU compared to GFP-expressing controls (**Fig. 4B,C**, multiple unpaired t-tests with Holm-Šídák’s correction, p > 0.05, N = 11 mice/group). These results suggest that increased MCU expression does not enhance respiration by increasing the expression levels of the assessed complex subunits. To confirm this effect was not due to differences in mitochondrial content between groups, we compared VDAC expression relative to total protein in mitochondrial fractions. We found no significant difference in the mitochondrial content of MCU OE and GFP-expressing mitochondrial fractions (**S.4**, avg. rel. density ±SEM, GFP: 1.0 ±0.14; MCU OE: 1.0 ±0.12; two-tailed unpaired t–test, p >0.05, N = 11 mice/group).

**Figure 4.**
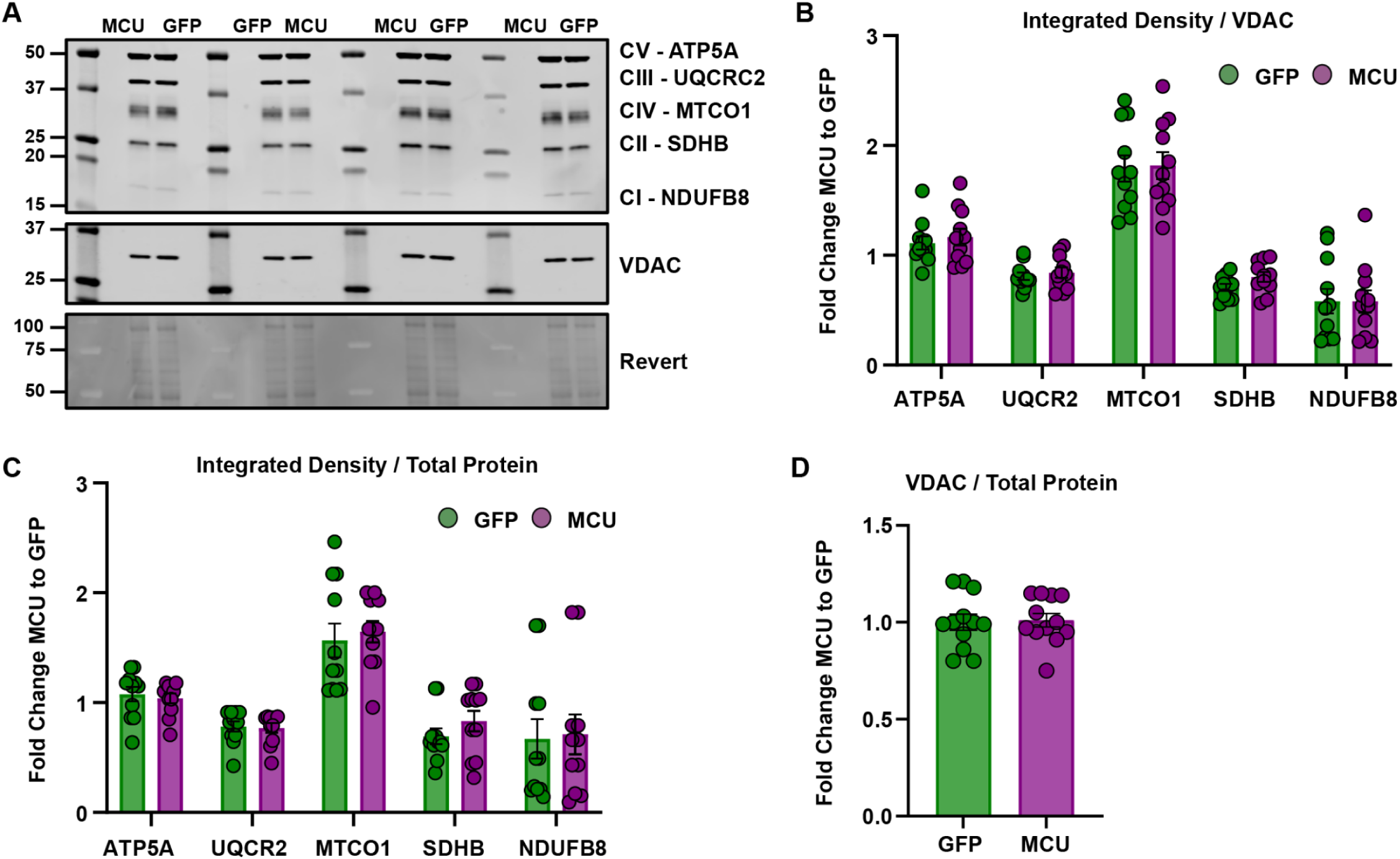
MCU overexpression does not increase the expression of ETC complex subunits. A. Representative immunoblots using antibodies against ETC complex I subunit NDUFB8, complex II subunit SDHB, complex III subunit UQCR2, complex IV subunit MTCO1, and complex V subunit ATP5A in mitochondria isolated from hippocampi overexpressing GFP or MCU. VDAC was used as a mitochondrial loading control. Full uncropped membranes and replicate blots corresponding to representative images are shown in the supplementary materials (**S.8,9,10**). B. Quantification shows the integrated density of ETC complex I-V subunits relative to the loading control VDAC. Densitometry was reported as a fold change relative to GFP controls (avg. F.C. ATP5A to GFP ±SEM, GFP: 1.11±0.06; MCU OE: 1.17±0.08; avg. F.C. UQCR2 to GFP ±SEM, GFP: 0.81±0.03; MCU OE: 0.84 ±0.05; avg. F.C. MTCO1 to GFP ±SEM, GFP: 1.79 ±0.12; MCU OE: 1.82±0.12; avg. F.C. SDHB to GFP ±SEM, GFP: 0.71 ±0.03; MCU OE: 0.80 ±0.04; avg. F.C. NDUFB8 to GFP ±SEM, GFP: 0.58 ±0.11; MCU OE: 0.58 ±0.10; multiple unpaired t-tests; all comparisons p > 0.05, corrected Holm-Ṧídák’s tests, N = 11 mice/group). C. Quantification shows the integrated density of ETC complex I-V subunits relative to total protein. Densitometry was reported as a fold change relative to GFP controls (avg. F.C. ATP5A to GFP ±SEM, GFP: 1.08 ±0.07; MCU OE:1.04 ±0.05; avg. F.C. UQCR2 to GFP ±SEM, GFP: 0.78 ±0.05; MCU OE:0.77 ±0.04; avg. F.C. MTCO1 to GFP ±SEM, GFP: 1.57 ±0.16; MCU OE: 1.65 ±0.10; avg. F.C. SDHB to GFP ±SEM, GFP: 0.69 ±0.07; MCU OE: 0.83 ±0.09; avg. F.C. NDUFB8 to GFP ±SEM, GFP: 0.67 ±0.18; MCU OE: 0.71 ±0.18; multiple unpaired t-tests; SDHB p = 0.76, all other comparisons p = 0.99, corrected Holm-Ṧídák’s tests, N = 11 mice/group). D. Quantification shows the integrated density of VDAC relative to total protein from mitochondrial fractions. Densitometry was reported as a fold change relative to GFP controls per cohort (avg. rel. density ±SEM, GFP: 1.04 ±0.14; MCU OE: 1.04 ±0.12; two-tailed unpaired t–test, p >0.05, N = 11 mice/group).

## 3 DISCUSSION

In this study, we found that increased expression of the MCU pore in hippocampal neurons lowers the MICU1:MCU ratio and increases the rate of MCU-dependent mitochondrial calcium uptake compared to controls, but does not affect calcium retention capacity (CRC) or sensitivity to mitochondrial calcium overload. Overexpression of MCU also resulted in significant increases in mitochondrial oxidative phosphorylation and improved electron transport chain efficiency under intense ATP regeneration demand. Together, these data indicate that MCU overexpression is sufficient to enhance mitochondrial bioenergetics in proportion to bioenergetic demand in isolated hippocampal mitochondria. However, overexpressing MCU was not accompanied by increased ETC complex protein expression, suggesting that enrichment of MCU does not upregulate mitochondrial biogenesis, but may instead fine-tune bioenergetics by increasing the efficiency of ATP synthesis, perhaps through matrix remodeling. Overexpressing MCU could drive adaptations in matrix calcium handling to increase oxidative phosphorylation in neurons with greater activity demand. These findings may broadly apply to mitochondrial heterogeneity and/or cell-type-specific adaptations to meet differential energy demands across neuronal cell types. While MCU overexpression was specific to hippocampal neurons (excitatory and inhibitory), we isolated mitochondria from the entire hippocampus, so our crude fractions included mitochondria from other cells (e.g., glia, traversing axons, vasculature) that were not transduced. Thus, there is the potential that our results could be attributable to (or confounded by) non-cell autonomous effects from these other cell types or that their mitochondria diminish the effect sizes from MCU overexpressing neurons. Nevertheless, the number of transduced neurons and/or the effect sizes were sufficient to detect differences from the total hippocampus. With advancements in cell-specific mitochondrial isolations (Fecher *et al*., 2019; de Mello *et al*., 2023), it is technically feasible for future studies to manipulate and isolate mitochondria from defined cell types to directly assess how differences in mitochondrial protein composition impact mitochondrial function.

### 3.1 MCU overexpression increases the rate of mitochondrial calcium uptake in hippocampal neurons

Enrichment of MCU was previously linked to enhanced calcium handling in cell-specific mitochondria isolated from the mouse brain. Similar to hippocampal neurons, different cell types in the mouse cerebellum differentially express MCU complex proteins. Granule cell mitochondria express significantly more MCU complex proteins than Purkinje cells and astrocytes, and showed a greater difference in mitochondrial calcium using area under the curve analyses relative to Ru360 (ΔAUC/ΔAUC_Ru360_), suggesting enhanced calcium uptake capacity (Fecher *et al*., 2019). In addition, mouse primary cortical neurons overexpressing MCU showed a greater ratiometric increase (ΔR/R0) in KCl-induced (Granatiero *et al*., 2019) or NMDA-evoked (Qiu *et al*., 2013) mitochondrial calcium levels compared to control neurons, also suggesting enhanced calcium uptake capacity. On the other hand, shRNA-mediated silencing of MCU in hippocampal neurons reduced NMDA-evoked mitochondrial calcium uptake compared to controls (Qiu *et al*., 2013). Genetic deletion of one or both MCU alleles in cerebellar granule cells resulted in a dose-dependent suppression of calcium uptake in isolated mitochondria (Fecher *et al*., 2019). In the acute hippocampal slice, MCU global haploinsufficiency reduced electrical- and glutamate-stimulated mitochondrial calcium uptake in mossy fiber axons (Devine *et al*., 2022). Collectively, these studies provide evidence that endogenous enrichment or exogenous expression of the MCU pore in neurons is sufficient to enhance mitochondrial calcium uptake, whereas mitochondria with diminished MCU expression exhibit reduced calcium uptake.

One factor that could explain how overexpressing MCU increases the rate of calcium uptake is a stoichiometric shift in the composition of MCU complex proteins. The ratio of MICU1 to MCU varies across tissue types (Márkus *et al*., 2016; Paillard *et al*., 2017; Farris *et al*., 2019), and this asymmetry regulates pore conductivity (Paillard *et al*., 2017). Across hippocampal subregions, CA2 neurons have the most *Mcu* expression and the lowest Micu1:Mcu ratio, whereas CA1 neurons have the least *Mcu* and the highest Micu1:Mcu ratio, suggesting that each subregion has a different mitochondrial calcium uptake profile (Márkus *et al*., 2016; Farris *et al*., 2019). In our study, MCU overexpression reduced the MICU1:MCU ratio, thereby increasing the rate of mitochondrial calcium uptake relative to GFP-expressing controls. In zebrafish, MCU overexpression in photoreceptors led to decreased MICU3 levels, the neuron-specific MCU enhancer, as an adaptive response to limit calcium uptake (Hutto *et al*., 2020). In cardiomyocytes, increasing MICU1 levels, and therefore the MICU1:MCU ratio, increased the calcium threshold of the MCU pore (Paillard *et al*., 2017). Our findings align with prior evidence that altering the MICU1:MCU expression ratio can modulate mitochondrial calcium uptake. However, our data cannot rule out compensatory responses by the other regulatory subunits. Though expression of Micu3 and the dominant-negative regulator Mcub is very low in the hippocampus, and only Mcu, Micu1, and the scaffolding protein Mcur1 show differential expression across hippocampal subregions (Farris *et al*., 2019). Therefore, overexpressing MCU most likely increases the availability of MICU1-free MCU, thereby lowering the channel threshold for calcium and enhancing calcium uptake rates. Follow-up studies at submicromolar calcium concentrations are needed to determine whether the increased uptake rate reflects a lower uptake threshold (Paillard *et al*., 2017).

### 3.2 MCU overexpression does not enhance the capacity for calcium or sensitivity to calcium overload

Excessive matrix calcium can trigger mPTP opening, leading to mitochondrial membrane rupture, the release of mitochondrial contents, and the activation of cell death pathways (Bernardi *et al*., 2023). Despite MCU overexpression, CRC and sensitivity to calcium overload remained unchanged in MCU OE and GFP mitochondria, which is at odds with several previous studies. Mouse cortical neurons overexpressing MCU *in vitro* showed greater ratiometric (ΔR/R0) decay in tetramethylrhodamine methyl ester (TMRM) fluorescence, reflecting a loss of membrane potential resulting from excitotoxic calcium overload from glutamate (Granatiero *et al*., 2019) or NMDA (Qiu *et al*., 2013). This aligns with studies in the mouse cortex, which show that increased MCU pore expression *in vivo* was concurrent with gliosis and cell death (Granatiero *et al*., 2019). However, we observed that careful titration of MCU overexpression in the hippocampus did not lead to increased gliosis or cell death (Pannoni *et al*., 2023). Alternatively, intact hippocampal circuits may handle mitochondrial calcium differently from cortical neurons. Consistent with this, in acute brain slices, hippocampal CA1 neurons took up more mitochondrial calcium at lower action potential frequencies than cortical neurons, which exhibited slower decay, suggesting region-specific differences in the coupling between action potential firing and MCU-dependent calcium uptake (Groten and MacVicar, 2022). These findings suggest that different cell types may be differentially equipped to handle mitochondrial calcium overload.

Alternatively, overexpressing MCU could have resulted in a baseline shift in mitochondrial calcium levels, thereby masking differences in the CRC. Hippocampal CA2 distal dendrites expressed more MCU and exhibited higher basal mitochondrial matrix calcium levels than proximal dendrites (Alsalman *et al*., 2025). Cultured mouse cortical neurons and adult zebrafish photoreceptors overexpressing MCU exhibited elevated basal matrix calcium levels (Qiu *et al*., 2013; Hutto *et al*., 2020). Elevated basal matrix calcium should reduce apparent capacity, causing mitochondria to reach calcium overload at similar time points despite underlying differences in CRC. Future studies should incorporate a mitochondrial matrix calcium-sensitive dye or sensor to help reconcile whether overexpressing MCU increases mitochondrial baseline calcium levels in isolated mitochondria. Compensatory changes in efflux could also explain why sensitivity to calcium overload remained unchanged in mitochondria overexpressing MCU. Increasing calcium efflux should lower matrix calcium levels and restore the threshold for mPTP activity (Bernardi *et al*., 2023). Restoring PKA-dependent phosphorylation of sodium-calcium-lithium exchanger (NCLX) in PINK1 KO dopaminergic neurons previously rescued mitochondrial calcium efflux deficiency and prevented mPTP-triggered cell death (Kostic *et al*., 2015). Our study cannot rule out changes in calcium efflux because we did not directly inhibit NCLX activity; future studies should use an NCLX inhibitor to test for compensatory efflux. Furthermore, measuring absorbance, or the loss of light scattering due to mitochondrial swelling in response to lower calcium concentrations, will help determine whether MCU overexpression increases sensitivity to mPTP activity. Clarifying the relationship between MCU differential expression and mPTP activity will be important for understanding how MCU may influence cell-type-specific vulnerabilities to calcium overload.

### 3.3 MCU overexpression enhances mitochondrial bioenergetics in hippocampal neurons

Overexpressing MCU in hippocampal neurons was sufficient to enhance complex I and II-stimulated oxidative phosphorylation and respiratory conductance, revealing a previously unrecognized role for the MCU pore in ETC efficiency in response to bioenergetic demand. This data suggests that neurons with low MICU1:MCU ratios may be more adept at responding to activity-driven changes in ATP demand. Overexpressing MCU for weeks could have led to a prolonged elevation in matrix calcium levels, thereby adapting mitochondria to higher demands and enabling more efficient ATP synthesis. Previously, it was shown that increasing calcium to isolated rat brain mitochondria resulted in a dose-dependent (500 or 1000 nmol Ca^2+^/ mg mitochondria) increase in complex I, but not complex II-stimulated basal oxygen consumption (Pandya, Nukala and Sullivan, 2013). Whereas ADP-stimulated and FCCP-uncoupled oxygen consumption declined steeply relative to controls with no calcium added (Pandya, Nukala and Sullivan, 2013). These data indicate that relatively high concentrations of acute calcium boost basal levels of respiration but inhibit maximal capacity for complex I ATP synthesis. Furthermore, calcium-dependent inhibition of oxidative phosphorylation was entirely independent of complex II (Pandya, Nukala and Sullivan, 2013). A similar effect was reported in isolated mitochondria from mouse liver, which exhibited increased respiratory control ratios (a measure of oxidative phosphorylation efficiency) under low-calcium complex I (8 nmol Ca^2+^ / mg mitochondria), but not complex II (22 nmol Ca^2+^ / mg mitochondria), was stimulated with low-calcium (Vilas-Boas *et al*., 2023). Furthermore, blocking MCU-dependent calcium uptake or suppressing NCLX-dependent efflux in hepatocytes prevented calcium-induced increases in oxygen consumption under these low-calcium conditions (Vilas-Boas *et al*., 2023). In contrast to these findings, we report that overexpressing MCU increases complex I- and II-stimulated maximal respiration and is sufficient to increase complex I- and II-linked respiratory conductance independently, under physiological ATP:ADP ratios. Overexpressing MCU could drive long-term adaptations in matrix calcium handling by maintaining a steady influx of calcium at low concentrations and accelerating TCA cycle flux, thereby increasing the respiratory efficiency of complexes I and II.

Stimulating TCA cycle turnover can increase the availability of substrates like succinate and accelerate succinate dehydrogenase activity, thereby increasing electron flux to complex II. This is not the first connection drawn between differential MCU expression and calcium-dependent enzyme activity. Zebrafish photoreceptors overexpressing MCU showed increased production of TCA intermediates with U-13C-glucose labeling and enhanced α-ketoglutarate dehydrogenase activity with U-13C-glutamine labeling, although this manipulation over time was maladaptive for mitochondrial quality control (Hutto *et al*., 2020). Alternatively, overexpressing MCU could boost complex-II stimulated oxidative phosphorylation in ways independent of matrix calcium influx. Mitochondria from HeLa cells formed MICU1/MICU2 heterodimers on the inner boundary membrane when exposed to high extramitochondrial calcium and stimulated with ATP (Cohen *et al*., 2025). Heterodimerized MICU1/MICU2 bound to and increased the activity of a glycerol-3-phosphate dehydrogenase (GPD2) /succinate dehydrogenase (complex II) metabolon, producing more FADH_2_ and boosting electron transfer to complex III (Cohen *et al*., 2025). Although endogenous levels are low in the hippocampus, overexpression of MCU over the course of weeks could have increased the ratio of MICU1:MICU2 expression and the heterodimerization of MICU1/MICU2 *in vivo*. As a result, GPD2/complex II metabolon activity could have increased, thereby boosting oxidative phosphorylation independently of MCU-dependent calcium uptake (Cohen *et al*., 2025). Moreover, the substantial increase in complex II-mediated respiration we have reported in mitochondria overexpressing MCU could be the summed effect of both adaptations, reflecting a steady increase in TCA flux and GPD2/complex II metabolon activity.

Another potential explanation for the increase in both complex I- and II-stimulated oxidative phosphorylation was our novel use of the creatine kinase clamp, which significantly improved our ability to probe respiration and substrate preference in neuronal mitochondria (Fisher-Wellman *et al*., 2018). Traditional mitochondrial stress tests are a powerful measure of maximal capacity at supra-physiological ADP levels, whereas the CK clamp enables more subtle control over ATP:ADP ratios and, therefore, energetic demand. In doing so, we could have captured interactions typically masked by saturating ADP levels. The development of higher-resolution respirometry to capture more subtle differences in activity-dependent ATP synthesis may be extremely helpful for understanding physiological changes in calcium handling and for uncovering how mitochondrial adaptations subserve bioenergetic efficiency. It is possible that, given the significant energetic burden on neurons, mitochondrial adaptations have evolved to handle mitochondrial calcium differently across neuron cell types. Similar approaches could help tease apart how mitochondrial adaptations evolve to support distinct neuronal cell types and circuit function.

### 3.3 MCU overexpression does not increase the expression of ETC complex subunits

In the present study, we modeled *in vivo* supply and demand using a creatine kinase clamp to assess the functional significance of MCU enrichment on oxidative phosphorylation. Elevated levels of MCU expression increased the efficiency of mitochondrial respiration at physiologically relevant ATP:ADP ratios, but we saw no difference in the expression levels of ETC complex subunits. This was not due to differences in overall mitochondrial content between groups, as evidenced by similar levels of the outer mitochondrial membrane protein VDAC. However, we examined only a small subset of the hundreds of ETC subunits and may have missed changes in other subunits we did not measure. An increase in respiration without a change in complex subunit expression could be indicative of ETC complex reorganizations or the formation of supercomplexes. Alternatively, enhanced respiration and calcium uptake could be a product of a more stable IMM, which in turn, enhances the stability of most major machinery in the matrix, including the MCU pore, ETC complexes I-V, and enables the formation of supercomplexes (Ghosh *et al*., 2020; Golla, Boyd and May, 2024; Bochkova *et al*., 2025; Emaus *et al*., 2025). Future studies investigating supercomplex architecture and mitochondrial ultrastructure can help resolve whether structural adaptations contribute to MCU-dependent enhancement of respiration.

Collectively, this work in isolated mitochondria establishes a foundation for dissecting how differential MCU expression tunes calcium handling and bioenergetic properties in intact circuits. Differential expression of MCU in neurons may be an important mechanism to ensure that mitochondrial calcium handling and ATP production are proportionate to the bioenergetic demands of individual cell types. Therefore, regions with mitochondrial heterogeneity, like the hippocampus (Cawley and Farris, 2026), may exhibit significant regional vulnerability to mitochondria-related neuropathologies. A better understanding of how mitochondrial heterogeneity supports diverse metabolic demands could aid in identifying cell types vulnerable to mitochondrial dysfunction and in developing novel ways to combat neurodegenerative disorders.

## 4 METHODS

### 4.1 Animals

Sexually naive adult (P50-100) male and female C57BL/6 background (C57BL/6J: B6, Jax #000664; C57BL/6NJ: B6N, Jax #005304; TdTomatofl/fl (Ai14(RCL-tdT)-D, Jax #007908, Amigo2-EGFP (Tg(Amigo2-EGFP)LW244Gsat/Mmucd, MMRRC#033018-UCD) mice were group-housed under a 12:12 light/dark cycle with access to food and water ad libitum. All procedures were approved by the Animal Care and Use Committee of Virginia Tech and were in accordance with the National Institutes of Health guidelines for care and use of animals.

### 4.2 Viral Infusion

Prior to surgery, mice were weighed and acclimated to the room before receiving an intraperitoneal (IP) injection of a ketamine/dexmedetomidine cocktail (ketamine, 50-100 mg/kg; dexmedetomidine, 0.375-0.5 mg/kg) for anesthesia. Mice were then placed on a heat pad, and an eye lubricant was applied before the hair around the surgical area was trimmed. The mouse was then placed on a stereotaxic apparatus with heat provided throughout the procedure. Mice received a subcutaneous injection of extended-release buprenorphine (3.25 mg/kg), and the area was disinfected with 3 cycles of Betadine and 70% ethanol before a longitudinal incision was made. Once the incision was made, the skull was cleaned, and bilateral burr holes were drilled targeting hippocampal CA1 and DG (−2.02 mm AP; +/- 1.50 mm ML; -1.55 mm DV). With a glass pipette and a syringe pump, 0.3µl of AAV9-hsyn1-mMCU-eGFP (Vector Biolabs, AAV-254662) or AAV9-hSyn1-eGFP (Vector Biolabs, VB1107) was infused at a rate of 50nL/min. The experimenter waited 5 min before pulling out the glass pipette. The mouse was then removed from the stereotaxic apparatus, and the incision was closed using surgical glue. Mice were administered an intraperitoneal (IP) injection of antisedan (1-2 mg/kg) and a subcutaneous injection of extended-release buprenorphine (3.25 mg/kg). Mice were then allowed to recover on a heating pad until they could ambulate. Functional assays were performed 16-21 days after surgery.

### 4.3 Mitochondrial Isolation

Differential centrifugation was used to isolate mitochondria from hippocampal brain tissue as previously described (Fisher-Wellman *et al*., 2018; Montalvo *et al*., 2026). Hippocampal tissue was homogenized with a Polytron at 50% maximum speed in ice-cold Mitochondrial Isolation Buffer (pH = 7.4; 2mg/mL bovine serum albumin, sucrose (70 mM), Mannitol (210 mM), HEPES (5 mM), EGTA (1 mM)). Tissue homogenates were centrifuged at 850 x g for 10 minutes at 4 °C. The tissue supernatant was centrifuged at 9000 ×g for 10 minutes at 4°C. The supernatant was then removed for western blot analysis of the cytosolic fractions. Mitochondrial pellets were suspended in 75 μL of Calcium Uptake Buffer (CUB; pH = 7.4, KCl (120 mM; Sigma P4504), MOPS (5 mM; Sigma), KH2PO4 (5 mM; Sigma P0662)). Protein content was determined using the Bradford Assay.

### 4.4 Western Blot and Electron Transport Chain Protein Expression

7.5 µg of mouse hippocampal cytosolic and mitochondrial fractions were resolved using 4-20% polyacrylamide gel electrophoresis (PAGE) under denaturing conditions (Criterion Midi Protein Gel, Bio-Rad). Following transfer to nitrocellulose membranes, anti-MCU (ProteinTech-26312-1-AP), -MICU1 (Sigma-HPA037480), -VDAC (Abcam-ab306581), and -AKT (Cell Signaling-9272) antibodies were used at a dilution of 1:2000 in 3% bovine serum albumin (BSA; Fisher Scientific) in TBS-T. Total OXPHOS (Abcam-110413) was used at a 1:500 dilution in 1% milk in PBS. Signals were detected using anti-rabbit 680 or anti-mouse 800 secondary antibodies at a dilution of 1:10,000 in 3% BSA in TBS-T or 1% milk in PBS, followed by washing and imaging using the Odyssey CLx Imaging System (LI-COR). Quantification was performed using densitometry. Revert^TM^ 700 Total Protein Stain (LI-COR) was used to detect and quantify total protein per lane on nitrocellulose membranes.

### 4.5 Calcium Retention Capacity assay

Mitochondrial calcium retention capacity assays were adapted from previous methods (Parks, Murphy and Liu, 2018; Fecher *et al*., 2019; Mendoza and Karch, 2022) and optimized for mitochondrial isolations from nervous tissues. The Horiba Scientific QuantaMaster 4000 series from Photon Technology International fluorometer was used to measure the fluorescence intensity of the cell-impermeable Ca2+ indicator Calcium Green-5 N (CG5N; 0.3 mM, excitation at 506 nm, emission at 532 nm, 25 °C, stirring at 600 rpm) upon stimulation with calcium chloride. 125 μg of mitochondria in CUB were loaded into a clean 875 μL Quartz cuvette (Fluoro-Open Top; Nova Biotech Qs-610) to a final volume of 600 μL and supplemented with Malate (5 mM; Sigma M7397), glutamate (10 mM; Sigma G5889), Ru360 (0.02 mM), or Cyclosporin A (100 μM). Cuvettes were incubated for 15 minutes under stirring before obtaining baseline measurements for background subtraction.

Sequential boluses of calcium chloride (Sigma 21115) were manually added to each cuvette every 7 minutes until fluorescence plateaued. Mitochondria were present at 125 µg per 600 µL (0.125 mg/mL), resulting in a Ca²⁺ load of 48 nmol Ca²⁺ per mg mitochondrial protein per pulse. At the plateau, an additional saturating dose of calcium chloride (1 M) was added to determine maximum fluorescence, followed by 0.5 M of the calcium chelator EGTA to determine minimum fluorescence. Each sample was paired with a Ru360 control under identical conditions and used to create a standard curve to assess calcium concentrations, calcium uptake rate, and retention capacity. The slope of peak Calcium Green™-5N intensity was used to extrapolate the rate and capacity of calcium uptake per mg of mitochondria in the absence and presence of Ru360.

### 4.6 Calcium Overload Assay

To investigate sensitivity to calcium overload, we adapted our CRC assay methods to measure increases in Calcium Green-5N (CG5N) fluorescence intensity resulting from elevated mitochondrial calcium release, serving as a proxy for swelling. Therefore, swelling is not directly quantified in these experiments, and the state of permeability transition was inferred from increases in CG5N fluorescence intensity, indicative of elevated free extramitochondrial calcium concentrations, following maximal calcium additions. Mitochondria were isolated and resuspended in CUB as previously described for CRC assays. Cuvettes with calcium swelling buffer (CSB: pH = 7.4, KCl (120 mM; Sigma P4504), MOPS (5 mM; SigmaM3183), KH2PO4 (5 mM; Sigma P0662)), Tris HCl (10 mM SigmaT1503) and energizing substrates malate (5 mM; Sigma M7397), glutamate (10 mM; Sigma G5889) were incubated for at least 7 minutes under stirring before obtaining baseline measurements for background subtraction.

Calcium Green-5 N (CG5N; 0.3 mM), 125 μg of isolated mitochondria, and Ru360 (0.02 mM) or Cyclosporin A (100 μM) were added to CSB to a final volume of 600 μL. MCU pore blocker Ru360 confirmed that mitochondrial calcium uptake, measured directly by CaG5N intensity loss, was due to MCU-dependent mitochondrial calcium uptake. The pharmacological inhibitor Cyclosporin A was used to block mPTP and confirm that increases in extramitochondrial calcium levels depended on mPTP activation. Calcium overload was achieved either sequentially or with a single large pulse of calcium chloride (Sigma 21115), which was manually added to each cuvette. CG5N traces were normalized using ΔF/F₀, where ΔF is CG5N baseline fluorescence (F₀) subtracted from CG5N fluorescence (F) at any point in the first 1200 seconds, and dividing ΔF by F₀. Peak CG5N was quantified from maximum ΔF/F₀ values over 1200 seconds (CG5N, Ru360) or 1000 seconds (CsA).

### 4.7 Mitochondrial Respiration

High-resolution oxygen consumption measurements were taken from isolated mitochondria using the Oroboros Oxygraph-2K (Oroboros, Innsbruck, Austria), as previously described (Fisher-Wellman *et al*., 2018; Montalvo *et al*., 2026). The O2k was set to and remained at 37°C and 500rpm to begin daily calibration with buffer D ( K-MES Potassium Salt (105mM; Sigma M0895), KCl (30mM; Sigma P4504), EGTA (1mM; Sigma E4378), KH2PO4 (10mM; Sigma P0662), MgCl2-6H2O (5mM; Sigma M2670), Bovine Serum Albumin (0.05%; Sigma A6003), pH = 7.1) to assess R1 (air saturation, ∼220 µM O2) and closed chamber R0 (zero oxygen) with sodium hydrosulfide (Sigma S1256). Chambers were then washed with deionized (DI) water (5 minutes for five cycles) before beginning the experiments.

We implemented the creatine kinase clamp method to assess oxygen flux (*J*O_2_) as previously described (Fisher-Wellman *et al.,* 2018; Montalvo *et al.,* 2026). This method provides a discrete and comprehensive measure of mitochondrial function by controlling energy demand and energy charge. 600 µL of BufferD + 5mM creatine monohydrate (Sigma C0780) was added to begin assessment of *J*O_2_ (pmol/s). Following chamber closure, 50µg of isolated mitochondria from the mouse hippocampus was added. Two separate conditions were implemented to differentiate the contribution of complex I from complex II respiration. To elucidate this question, we assessed respiration using creatine kinase (20 mM; Sigma C3755) in Buffer D supplemented with complex I-linked substrates, malate (5 mM; Sigma M7397), glutamate (10 mM; Sigma G5889), and fresh potassium pyruvate (5 mM; Combi-Blocks QA-1116). Separately, complex II-linked respiration was evaluated in Buffer D supplemented with creatine kinase (20 mM), succinate (10 mM; Sigma 53674), and rotenone (0.01 mM; Sigma R8875) to inhibit complex I. Maximal respiration and subsequent phosphocreatine titrations were performed identically for both substrate conditions. Maximal *J*O_2_ (State 3) was driven with the addition of phosphocreatine (PCr) (1 mM, Sigma P1937; ΔG_ATP_ -12.94 kCal/mol) and ATP (5 mM; Sigma A2383). Cytochrome C (0.005 mM, Sigma 2506) was then used to assess mitochondrial integrity, with a threshold of a 20% increase in respiration; samples with an increase greater than 20% were removed from analysis, but none were required (S.7). Sequential PCr titrations were then added at 1mM (ΔG_ATP_ -13.38 kCal/mol), 2mM (ΔG_ATP_ -13.71 kCal/mol), 4mM (ΔG_ATP_ -14.12 kCal/mol), and 6mM (ΔG_ATP_ -14.45 kCal/mol).

Calculations of ΔG_ATP_ were performed using an online calculator (Fisher-Wellman *et al*., 2018) under the conditions of 37°C, 170mM ionic strength, 5mM creatine, 10mM phosphate, and pH 7.1. Plotting *J*O_2_ vs. ΔG_ATP_ across the linear force-flow relationship enables calculation of mitochondrial respiratory conductance. Conductance measures the inverse of resistance (1/R) of the mitochondrial proton motive force (Δp; voltage) in relation to Ohm’s law (I=V/R), where current (I) measures JH+. Higher conductance (slope of *J*O_2_ vs. ΔG_ATP_) is akin to less resistance within the electron transport chain and the ability to respond to the energetic demand ΔG_ATP_. In this regard, measures of conductance over linear ranges of ΔG_ATP_ (i.e., PCr titrations) mimic a tractable bioenergetic stress test or transition from rest to exercise, as demonstrated in Figure 3.

### 4.8 Tissue Processing and Immunofluorescence

Mice were given an overdose of sodium pentobarbital (150 mg/kg) and perfused with 15-20 mL 4% paraformaldehyde for 4 min. Brains were post-fixed in 4% paraformaldehyde for at least 24 hr before being sectioned coronally at 40 μm on a Leica vibratome (VT1000S). Sections were then immunostained with rabbit-anti-RFP (1:500, Rockland, Cat#604-401-379, RRID:AB_2209751), chicken-anti-GFP (1:2000, Abcam, Cat#ab13970, RRID:AB_300798), and mouse-anti-Calb1 (1:500, Invitrogen, Cat#MA5-24135, RRID:AB_2609695) were used to label CA12. Tissues underwent antigen retrieval by boiling free-floating sections for 4-5 min in 1 mL nanopure water, followed by a 15-min permeabilization step in 0.1% Tween20 in 1X PBS (PBST). All sections were blocked for a minimum of 1 hr in 5% normal goat serum (NGS) in PBST, then incubated in primary antibodies overnight (18-24 hr) at room temperature. After 3 washes in PBST, sections were placed in a second block in 5% NGS in PBST for a minimum of 30 min, then incubated with secondary antibodies for 2 hr at room temperature. The secondary antibodies were diluted at 1:500 in 5% NGS in PBST blocking solution and included the following from Invitrogen: goat-anti-rabbit-546 Cat#A11035, goat-anti-chicken-488 Cat#A11039, goat-anti-mouse-488 Cat#A21037. Sectioned then underwent 3 washes in PBST and a last wash in PBS before they were mounted and coverslipped with prolonged gold fluorescence media with 4’,6-diamidino-2-phenylindole (DAPI; Invitrogen, Cat# P36931).

### 4.9 Image Acquisition and Quantification

Images were acquired on an inverted Leica Thunder epifluorescence microscope at 20x/0.8 NA. All images were acquired at 16-bit and subjected to computational clearing before quantification in Fiji (ImageJ).

### 4.10 Transmission Electron Microscopy

Isolated mitochondria in CSB were recovered from calcium overload assays and centrifuged at 600 x g for 5 minutes at 4 °C. The resulting supernatant was discarded, and the mitochondrial pellet was resuspended in ice-cold Karnovsky buffer (0.1 M cacodylate buffer, pH 7.4, containing 2.5% glutaraldehyde and 1.5% paraformaldehyde) at 4 °C for 2 weeks. Fixed samples were washed twice in 0.1 M cacodylate buffer for 5 minutes and post-fixed in 1% osmium tetroxide containing potassium ferricyanide for 30 minutes. Samples were rinsed in distilled water (x2 washes) and stained with 2% (w/v) uranyl acetate prepared in 25% ethanol for 1 hour. Graded ethanol was used to dehydrate samples: 50% ethanol (5min), 70% ethanol (10 min), 90% ethanol (10 min), and 100% ethanol (10 min). Samples were incubated in 1:1 ethanol/acetone for 10 minutes, and then again in 100% acetone for an additional 10 minutes. Mitochondria were placed in 1:2 acetone/Epon for 24 hours, transferred to 1:4 acetone/Epon for 1 hour, then to 100% Epon for 2 hours twice, followed by a final 100% Epon for 24 hours, before embedding in Epon resin and polymerizing in a drying oven. Ultrathin sections of mitochondria isolations were imaged using a FEI Tecnai G2 Spirit Twin Transmission Electron Microscope at 120 kV. Images were acquired using a Gatan Rio CCD camera at a magnification of 6800x, with final dimensions of 3072 x 3072 pixels.

### 4.11 Statistical analyses

All statistical analyses were performed using GraphPad Prism 9, and significance was determined at an alpha level of 0.05.

## DECLARATIONS

### Funding

Research reported in this publication was supported by the National Institute of Mental Health under award R01MH124997 to S.F., and by startup funds to S.F. provided by Virginia Tech. The funders had no role in the design of the study, collection, analysis, and interpretation of data, and in writing the manuscript.

### Authors’ contributions

Conceptualization, SF; Methodology, MLC, RNM, MLW, LLT, SF; Formal Analysis, MLC, RNM, MLW, LLT,SF; Investigation, MLC, RNM, MLW, SF; Writing – Original Draft, MLC, RNM, MLW, SF; Writing – Review & Editing, MLC, RNM, MLW, SF; Visualization, MLC, SF; Supervision, ZY, JP, SF; Funding Acquisition, SF.

## Acknowledgements

We thank the members of the Farris lab for providing feedback and critically reading this manuscript. The authors acknowledge resources and support from the Virginia Tech animal care staff and Cellular and Molecular Imaging Core, part of the Fralin Biomedical Research Institute at VTC. This research would not be possible without the use of the FEI Tecnai G2 Spirit Twin Transmission Electron Microscope, and the helpful insight in sample preparation and instrument operation from Christie Lacy and Clare Dennison.

## Conflict of interest

The authors declare that they have no competing interests.

## Notes

### Competing Interest Statement

The authors have declared no competing interest.

