## Supplemental Fig. 1-14 for "Elevating levels of neuronal MCU in the hippocampus enhances mitochondrial calcium uptake and respiratory efficiency proportional to demand"

**Supplemental Figures**

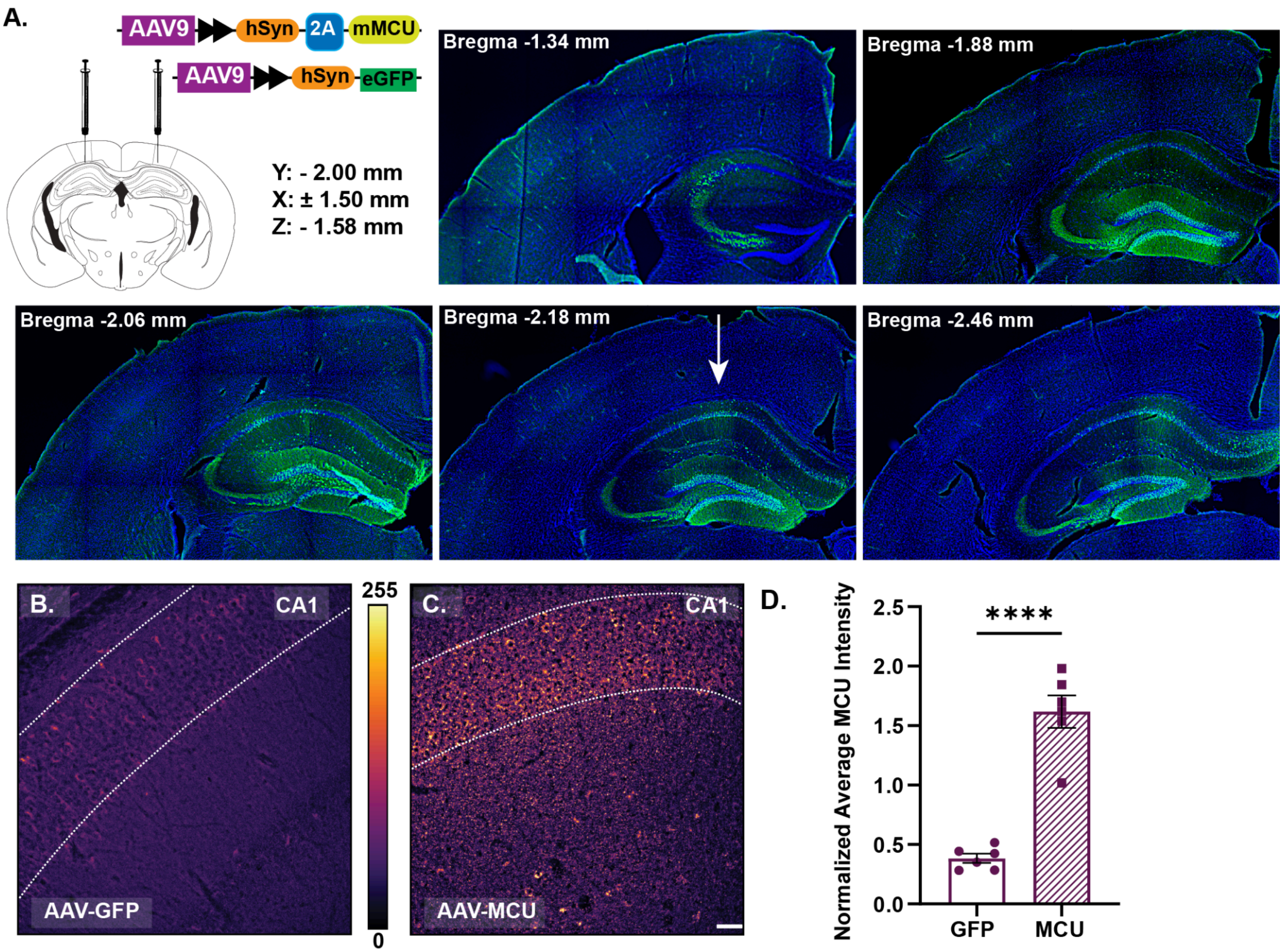


**Supplemental Figure 1:** *Validation of MCU overexpression in the mouse CA1 and DG hippocampal subregions.*

A) Representative 20X tile images of the MCU overexpression targeted to dorsal CA1 and DG hippocampal subregions. Coronal sections were immunostained with GFP to validate viral targeting and rostral-to-caudal spread. Images were used to determine ideal AAV titrations, limiting MCU enrichment to physiological levels to minimize off-target effects such as reactive gliosis or cell death. Scale = 300µm

B) Representative 20X image of MCU immunolabeling in the CA1 of mice expressing GFP.

C) Representative 20X image of MCU immunolabeling in the CA1 of mice overexpressing MCU.

D) Normalized average MCU immunofluorescence intensity was measured from ROIs that spanned all the CA1 laminae (stratum oriens, pyramidale, radiatum, and lacunosum moleculare) in GFP and MCU OE mice (avg. norm. int. ±SEM: GFP: 0.4 ±0.04; MCU OE: 1.6 ±0.1; two-tailed unpaired t-test; ****$p \leq 0.0001$, N = 6 mice/group). Quantitative images were acquired under identical parameters, and the background signal was subtracted before values were normalized to the cohort average. Scale = 50μm

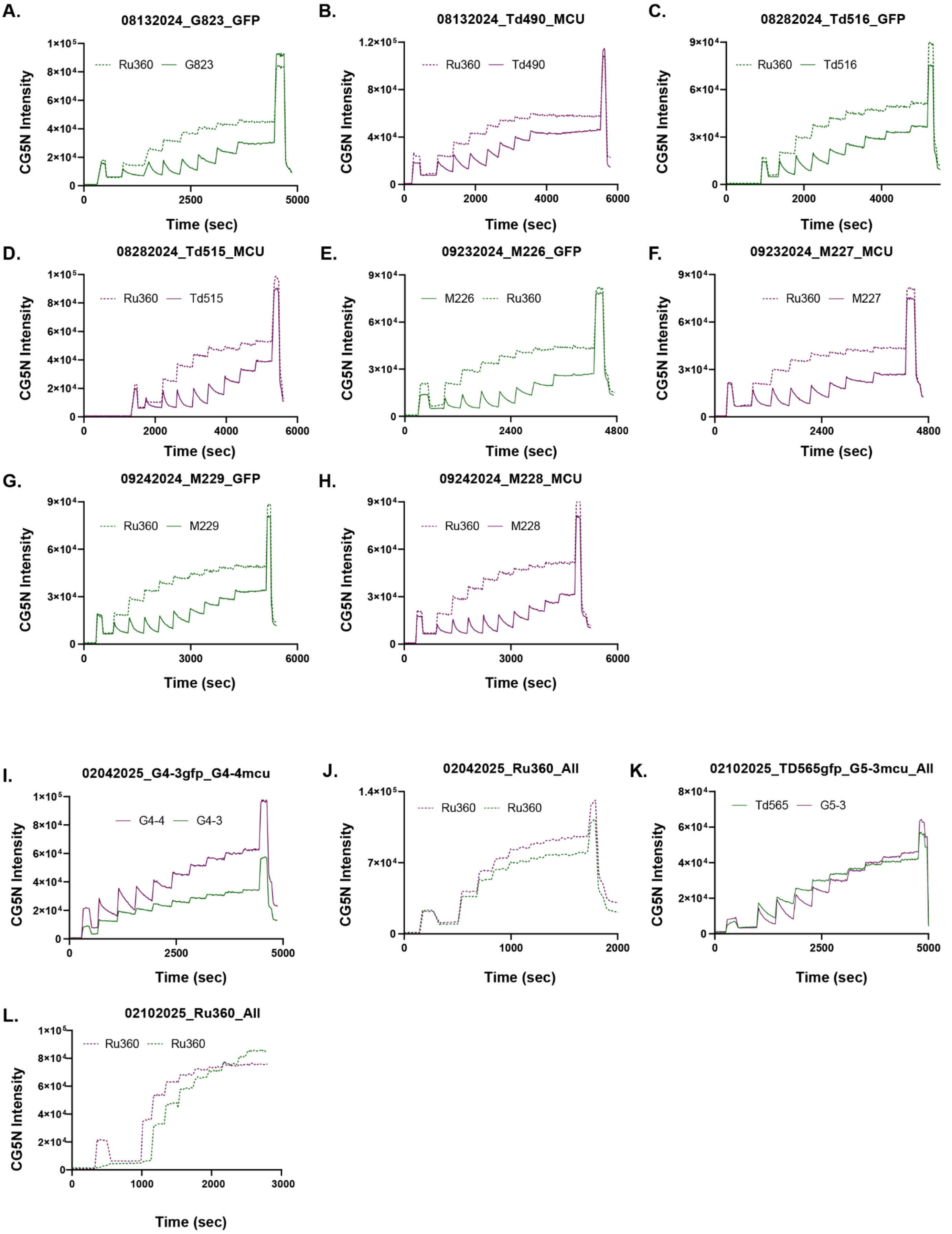
A.
08132024_G823_GFP
Ru360 G823
B.
08132024_Td490_MCU
Ru360 Td490
C.
08282024_Td516_GFP
Ru360 Td516
D.
08282024_Td515_MCU
Ru360 Td515
E.
09232024_M226_GFP
M226 Ru360
F.
09232024_M227_MCU
Ru360 M227
G.
09242024_M229_GFP
Ru360 M229
H.
09242024_M228_MCU
Ru360 M228
I.
02042025_G4-3gfp_G4-4mcu
G4-4 G4-3
J.
02042025_Ru360_All
Ru360 Ru360
K.
02102025_TD565gfp_G5-3mcu_All
Td565 G5-3
L.
02102025_Ru360_All
Ru360 Ru360
CG5N Intensity
Time (sec)

**Supplemental Figure 2.** *Unprocessed fluorescence traces from calcium retention capacity assays.*

A-L) Unprocessed traces for each CRC assay. Each panel shows traces from GFP (green; solid), MCU OE (magenta; solid), and Ru360 (dashed) controls. For a subset of CRC assays (I -L), runs were paired by experimental (MCU OE vs GFP) and control (GFP+RU360 vs MCU+RU360) groups. Standard curves were generated from RU360 controls immediately following experimental assays, but over a shorter period of time (<3000 seconds). See methods for more details.

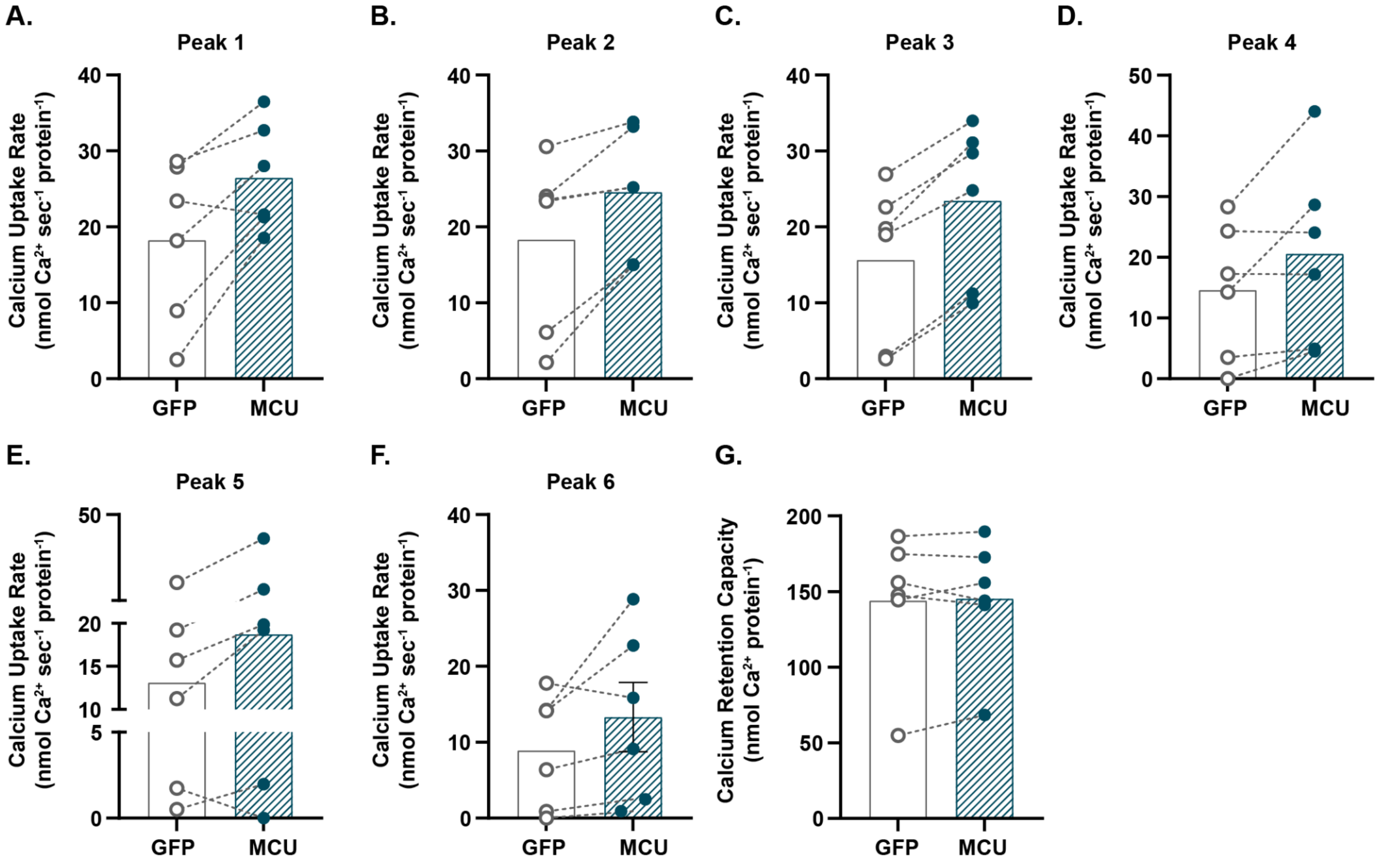


**Supplemental Figure 3.** *Calcium uptake rates and calcium retention capacity (CRC) pre-normalization by pair average.*

A-F) Quantification shows mitochondrial calcium uptake rates ($Ca^{2+}nmol \cdot s^{-1} \cdot mg^{-1}$) from isolated mitochondria expressing GFP or overexpressing MCU for each sequential $Ca^{2+}$ pulses (10 μmol) peak (1-6). Each point represents an independent biological replicate, and lines indicate GFP and MCU OE pairs. N = 6 mice/group.

G) Quantification shows the CRC ($Ca^{2+}nmol \cdot mg^{-1}$) from isolated mitochondria expressing GFP or overexpressing MCU in response to six sequential $Ca^{2+}$ pulses (Final: 60μmol $Ca^{2+}$). Each point represents an independent biological replicate, and lines indicate GFP and MCU OE pairs. N = 6 mice/group.

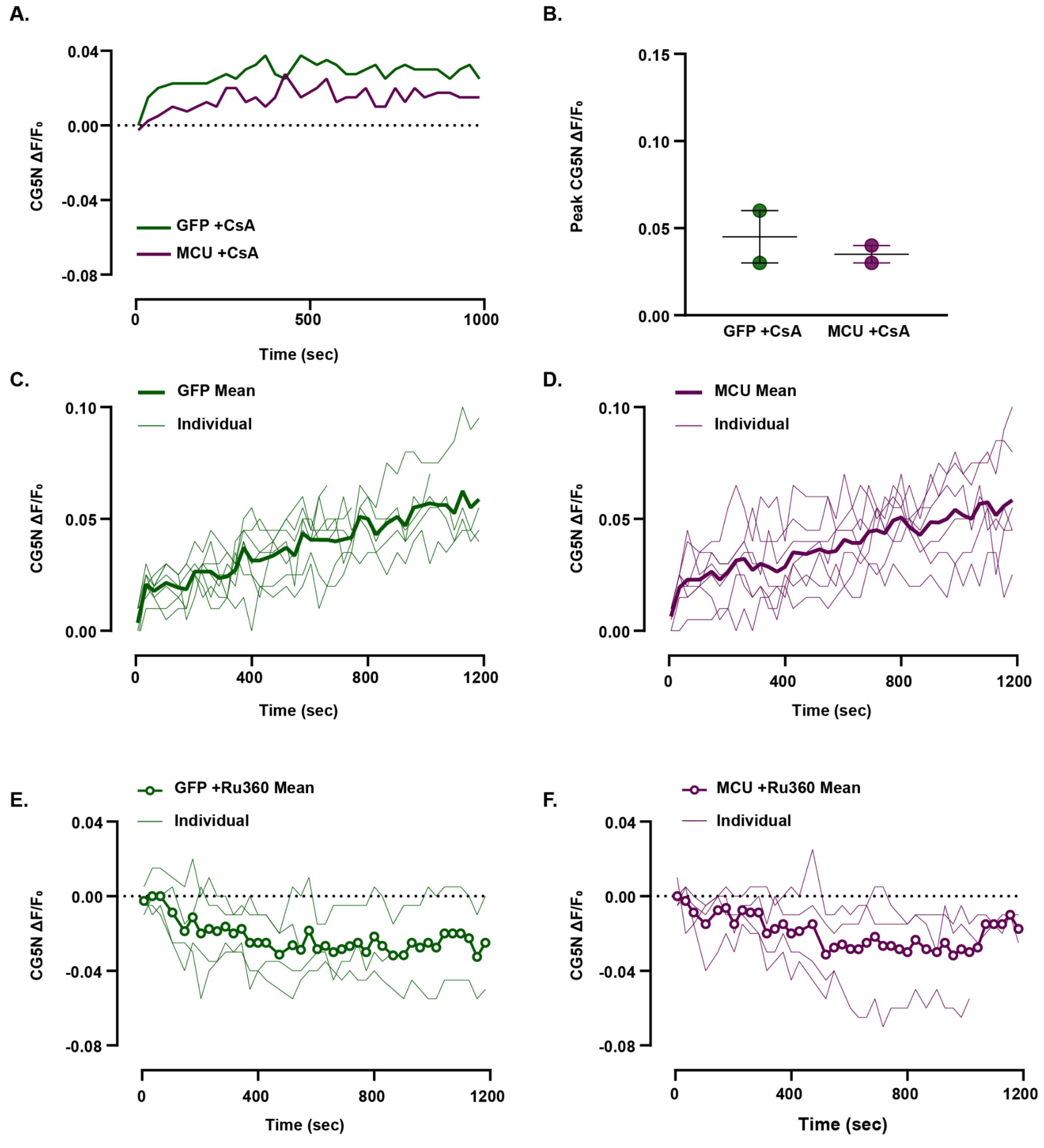


**Supplemental Figure 4.** *Representative individual CG5N intensity (ΔF/F0) traces and quantification*

A) Mean CG5N ΔF/F0 trace from MCU OE and GFP-expressing mitochondria in the presence of mPTP inhibitor CsA. Mean CG5N ΔF/F0 traces reflect the average trace per group recorded over 1000 seconds in response to large calcium pulses (range: 35-300μM). N = 2 mice/group.

B) Quantification for peak CG5N ΔF/F0 values from MCU OE and GFP-expressing mitochondria in the presence of mPTP inhibitor CsA. N = 2 mice/group.

C) GFP average (thick) and individual (thin) CG5N ΔF/F0 traces recorded over 1200 seconds in response to large calcium pulses (range: 35-300μM). N = 7 mice/group.

D) MCU OE average (thick) and individual (thin) CG5N ΔF/F0 traces recorded over 1200 seconds in response to large calcium pulses (range: 35-300μM). N = 7 mice/group.

E) GFP +Ru360 average (open circle) and individual (thin) CG5N ΔF/F0 traces recorded over 1200 seconds in response to large calcium pulses (range: 35-300μM). N = 4 mice/group.

F) MCU OE +Ru360 average (open circle) and individual (thin) CG5N ΔF/F0 traces recorded over 1200 seconds in response to large calcium pulses (range: 35-300μM). N = 4 mice/group.

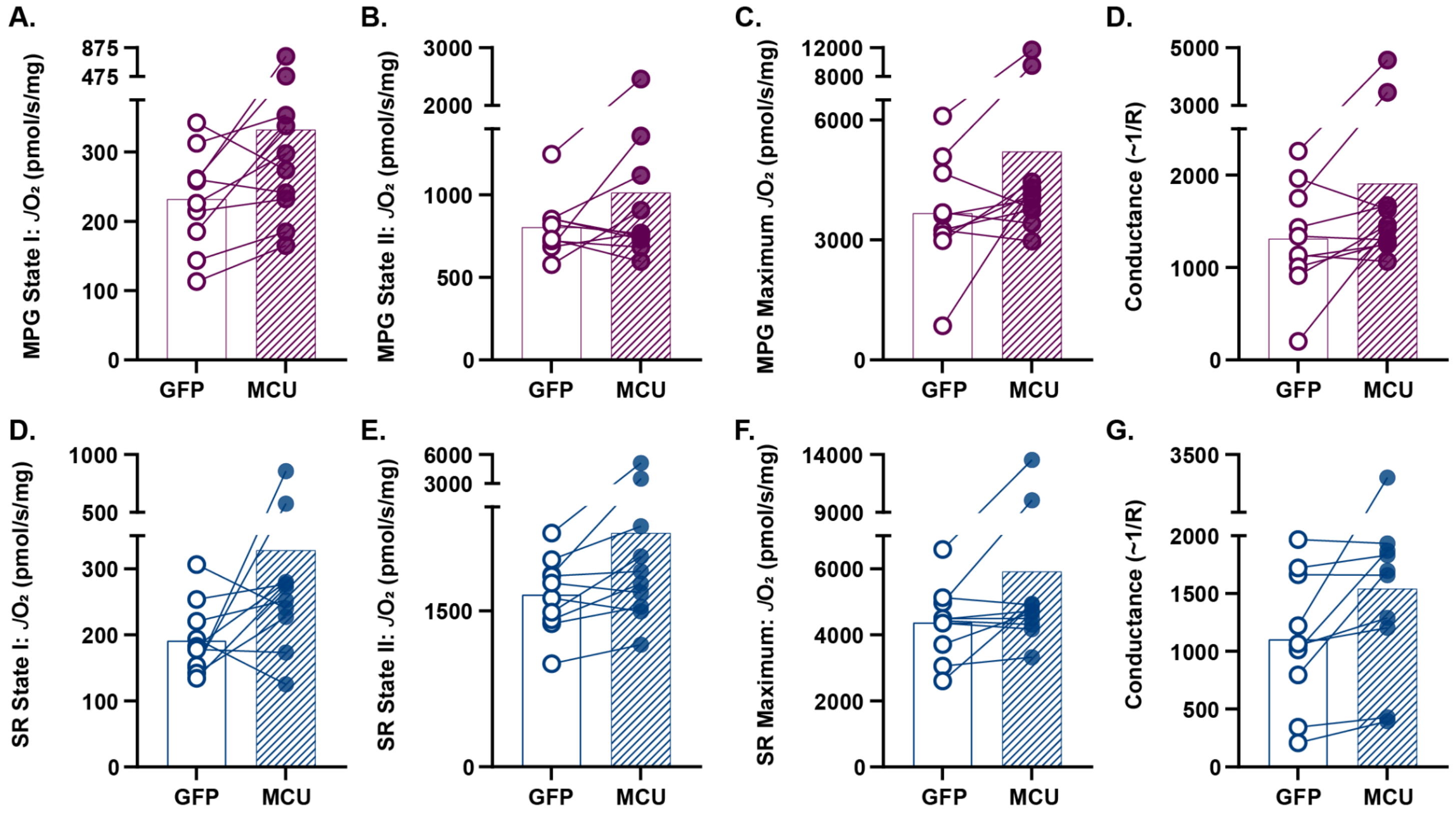


**Supplemental Figure 5.** *Raw, paired values for complex I and II-linked $JO_2$.*

A-D) Complex I-linked $JO_2$ (pmol $O_2 \cdot s^{-1} \cdot mg^{-1}$ mitochondrial protein) values for state I (mitochondria only), non-phosphorylating respiratory state (mitochondria, MPG), maximal $\Delta G_{ATP}$ -12.94 kCal/mol (mitochondria, MPG, ATP, phosphocreatine), and respiratory conductance (slope of $JO_2$ and $\Delta G_{ATP}$) measured in isolated mitochondria (50µg) expressing GFP or overexpressing MCU. N = 10 mice/group.

E-H) Complex II-linked $JO_2$ (pmol $O_2 \cdot s^{-1} \cdot mg^{-1}$ mitochondrial protein) values for state I (mitochondria only), non-phosphorylating respiratory state (mitochondria, SR), maximal $\Delta G_{ATP}$ -12.94 kCal/mol (mitochondria, SR, ATP, phosphocreatine), and respiratory conductance (slope of $JO_2$ and $\Delta G_{ATP}$) measured in isolated mitochondria (50µg) expressing GFP or overexpressing MCU. N = 10 mice/group.

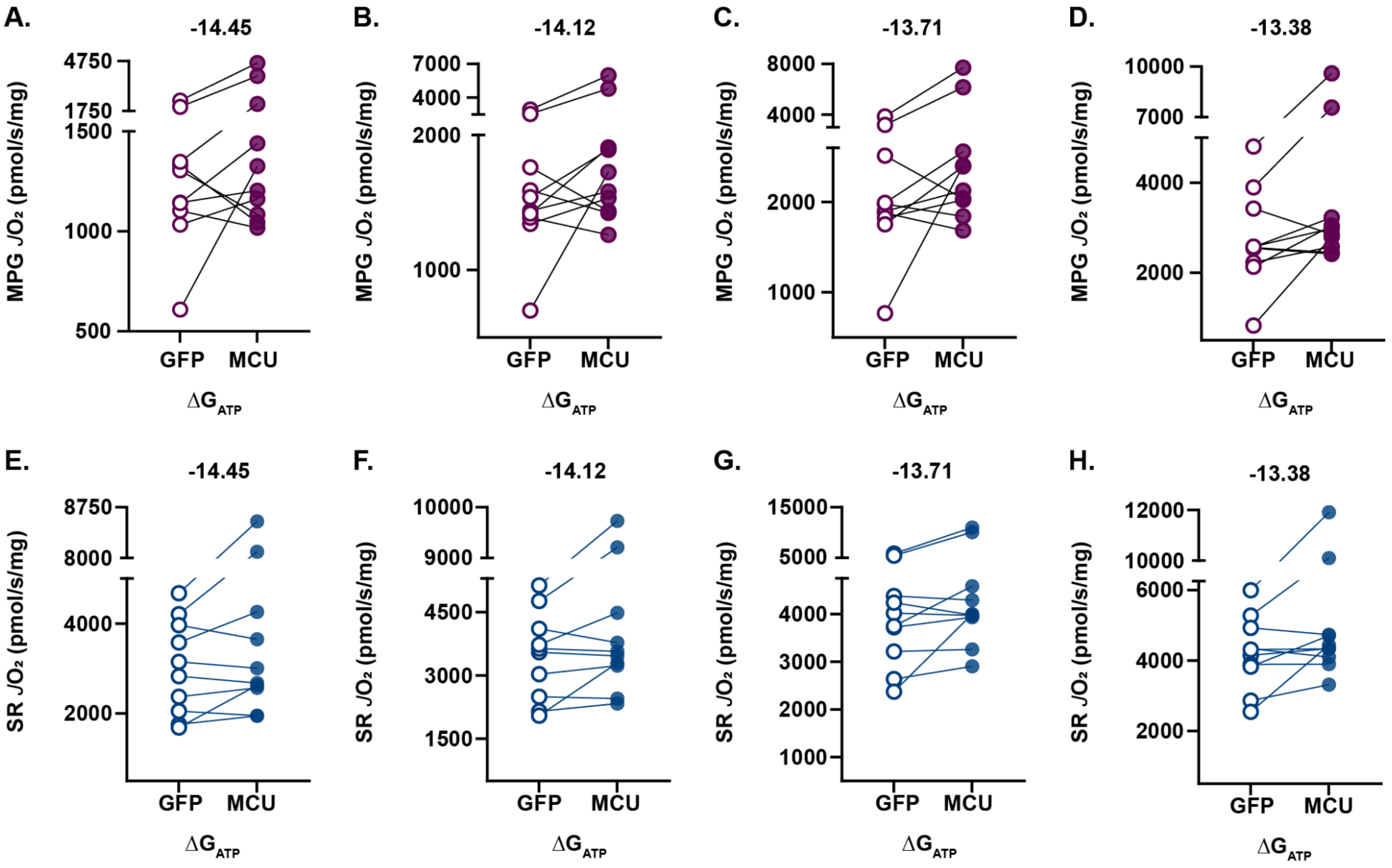


**Supplemental Figure 6.** *Raw, paired values for complex I and II-linked phosphocreatine titrations.* A-D) Complex I-linked $JO_2$ (pmol $O_2 \cdot s^{-1} \cdot mg^{-1}$ mitochondrial protein) in response to increasing $\Delta G_{ATP}$ measured by the phosphocreatine titration in isolated mitochondria expressing GFP or overexpressing MCU.

E-H) Complex II-linked $JO_2$ (pmol $O_2 \cdot s^{-1} \cdot mg^{-1}$ mitochondrial protein) in response to increasing $\Delta G_{ATP}$ measured by the phosphocreatine titration in isolated mitochondria expressing GFP or overexpressing MCU.

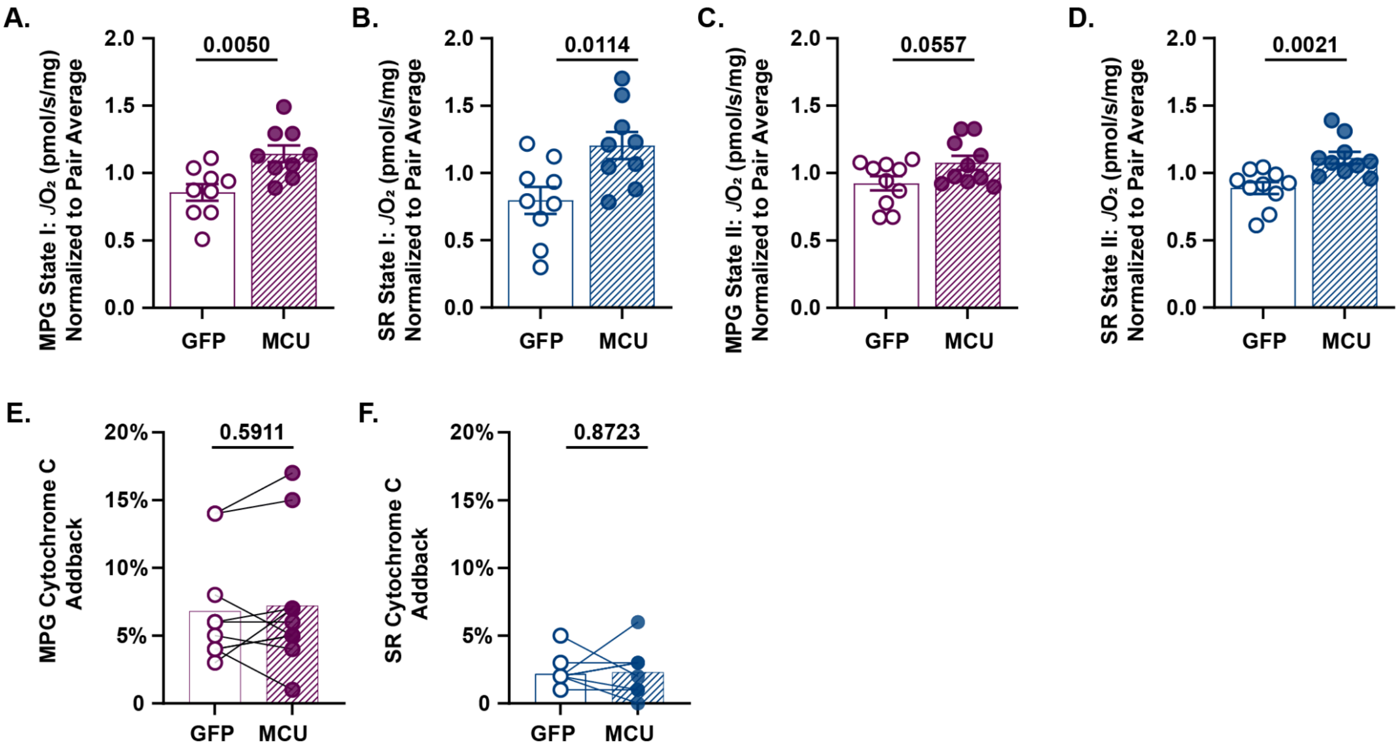


**Supplemental Figure 7.** *Normalized complex I and II-linked $JO_2$ values for state I, nonphosphorylating respiratory states, and cytochrome c addback.*

A. Normalized state I (mitochondria only) $JO_2$ (pmol $O_2 \cdot s^{-1} \cdot mg^{-1}$ mitochondrial protein) energized by MPG in isolated mitochondria (50µg) expressing GFP or overexpressing MCU (avg. norm. $JO_2$ ±SEM, GFP: 0.85 ±0.06; MCU OE: 1.10 ±0.06; two-tailed unpaired t-test; p = 0.005, N = 10 mice/group). Oxygen consumption rates are normalized by the average rate of each pair.

B. Normalized state I (mitochondria only) $JO_2$ (pmol $O_2 \cdot s^{-1} \cdot mg^{-1}$ mitochondrial protein) energized by SR in isolated mitochondria (50µg) expressing GFP or overexpressing MCU (avg. norm. $JO_2$ ±SEM, GFP: 0.82 ±0.09; MCU OE: 1.2 ±0.09; two-tailed unpaired t-test; p = 0.0114, N = 10 mice/group). Oxygen consumption rates are normalized by the average rate of each pair.

C. Normalized non-phosphorylating respiratory state $JO_2$ (pmol $O_2 \cdot s^{-1} \cdot mg^{-1}$ mitochondrial protein) energized by MPG in isolated mitochondria (50µg) expressing GFP or overexpressing MCU (avg. norm. $JO_2$ ±SEM, GFP: 0.92 ±0.05; MCU OE: 1.1 ±0.05; two-tailed unpaired t-test; p = 0.0557, N = mice/group). Oxygen consumption rates are normalized by the average rate of each pair.

D. Normalized non-phosphorylating respiratory state $JO_2$ (pmol $O_2 \cdot s^{-1} \cdot mg^{-1}$ mitochondrial protein) energized by SR in isolated mitochondria (50µg) expressing GFP or overexpressing MCU (avg. norm. $JO_2$ ±SEM, 0.89 ±0.04; MCU OE: 1.1 ±0.04; two-tailed unpaired t-test; p = 0.0021,

N = 10 mice/group). Oxygen consumption rates are normalized by the average rate of each pair.

E. Percent increase $JO_2$ % from cytochrome c add back in MPG stimulated isolated mitochondria expressing GFP or overexpressing MCU (avg. % $JO_2$ ±SEM, GFP: 6.8% ±1.3; MCU OE: 7.2% ± 1.6; two-tailed unpaired t-test; p = 0.5911, N = 10 mice/group).

F. Percent increase $JO_2$ % from cytochrome c add back in SR stimulated isolated mitochondria expressing GFP or overexpressing MCU (avg. % $JO_2$ ±SEM, GFP: 2.2% ±0.36; MCU OE: 2.3% ±0.54; two-tailed unpaired t-test; p = 0.8723, N = 10 mice/group).

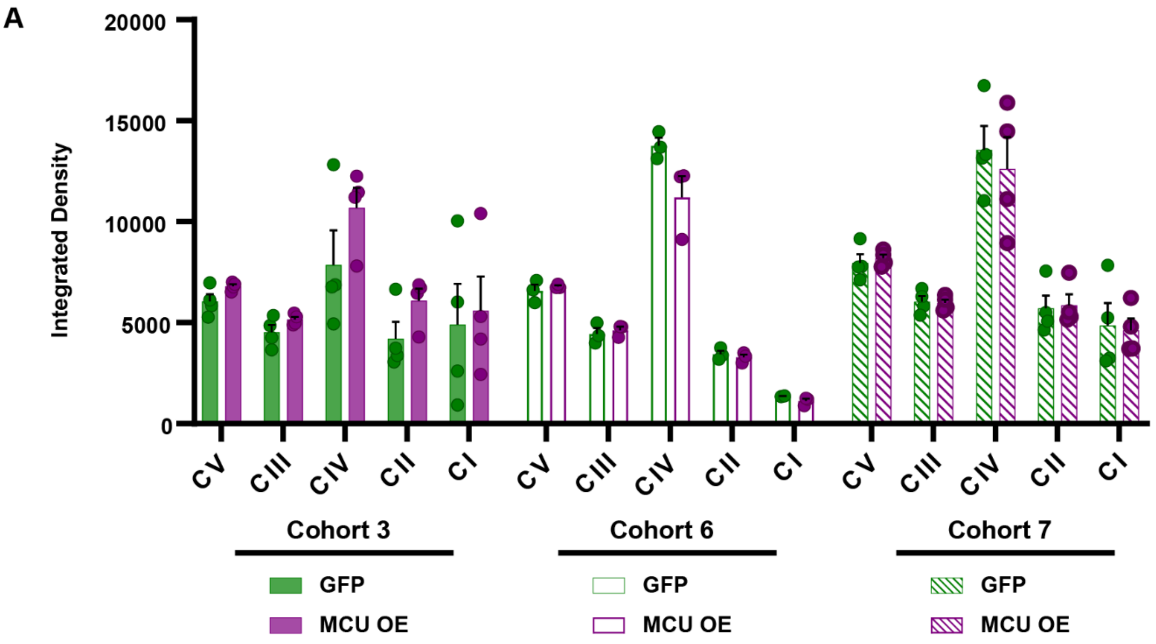


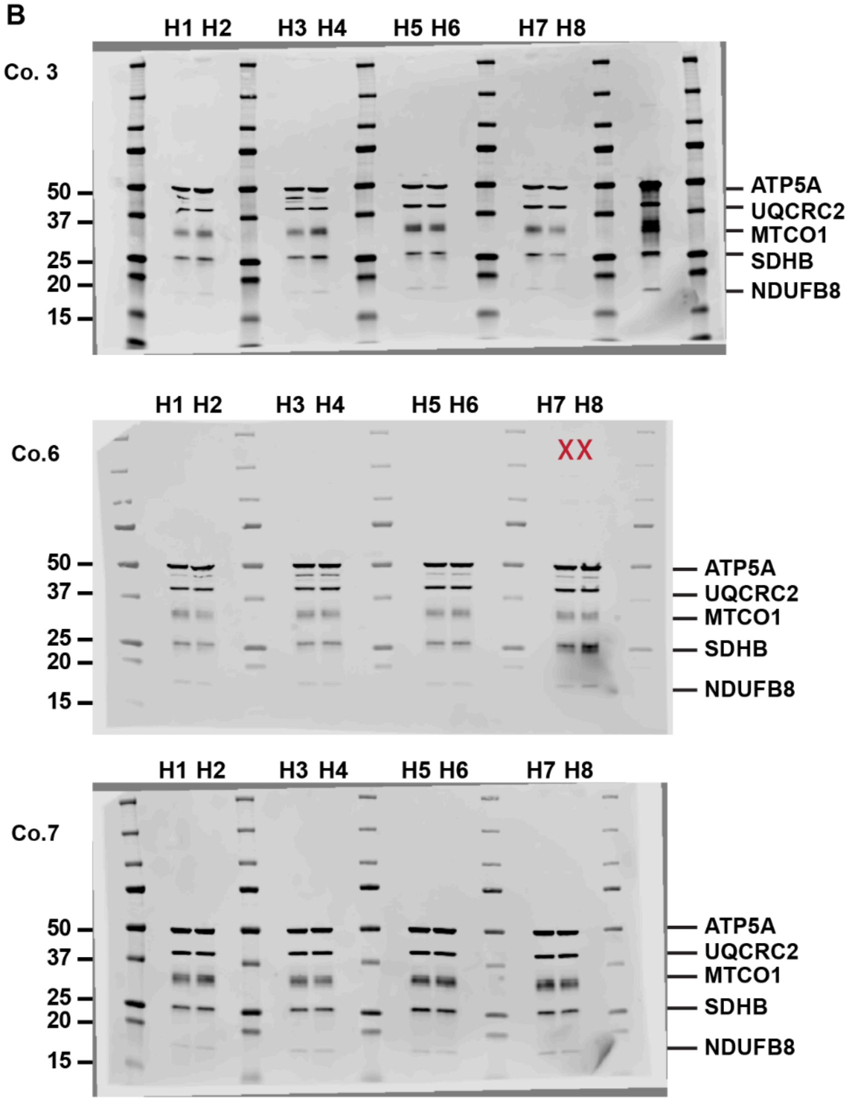


C

Cohort 3

| Lane | Ex. ID |
|---|---|
| H1 | G4 -3 |
| H2 | G4 -4 |
| H3 | Td565 |
| H4 | G5-3 |
| H5 | B008 |
| H6 | B005 |
| H7 | B007 |
| H8 | B006 |

Cohort 6

| Lane | Ex. ID |
|---|---|
| H1 | B036 |
| H2 | B026 |
| H3 | B035 |
| H4 | B038 |
| H5 | SS5 |
| H6 | SS4 |
| H7 | Excluded |
| H8 | Excluded |

Cohort 7

| Lane | Ex. ID |
|---|---|
| H1 | B070 |
| H2 | B071 |
| H3 | B074 |
| H4 | B072 |
| H5 | B080 |
| H6 | B081 |
| H7 | B085 |
| H8 | B086 |

**Supplemental Figure 8.** *Total OXPHOS integrated density across cohorts*

A. Quantification shows the integrated density of ETC complex I-V subunits from mitochondrial fractions overexpressing GFP or MCU across cohorts.

B. Uncropped immunoblots stained with Total OXPHOS antibodies against ATP5A, UGCRC2, MTCO1, SDHB, and NDUFB8 in mitochondrial (mito) fractions isolated from hippocampi overexpressing GFP or MCU. Densitometry was used to quantify the total band signal for each ETC complex subunit, which was then normalized to the corresponding VDAC loading control. Red X denotes samples excluded from the study.

C. Key provided for each experimental ID across cohorts.

**A**

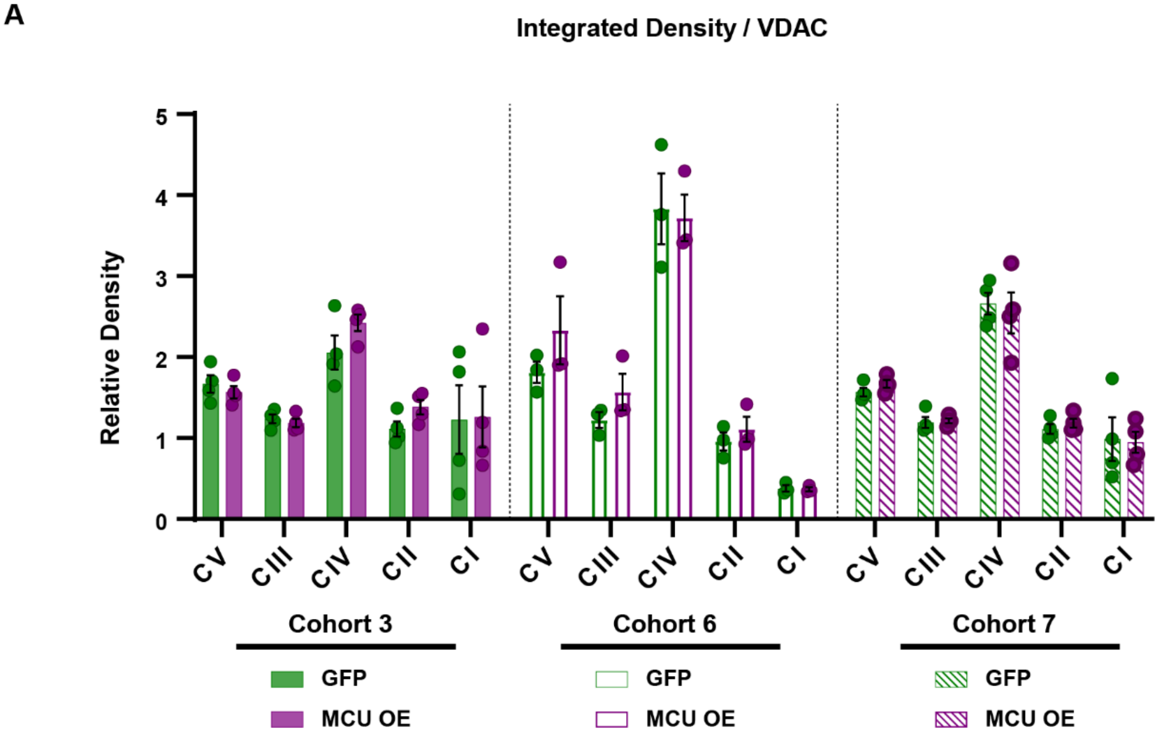


**B**

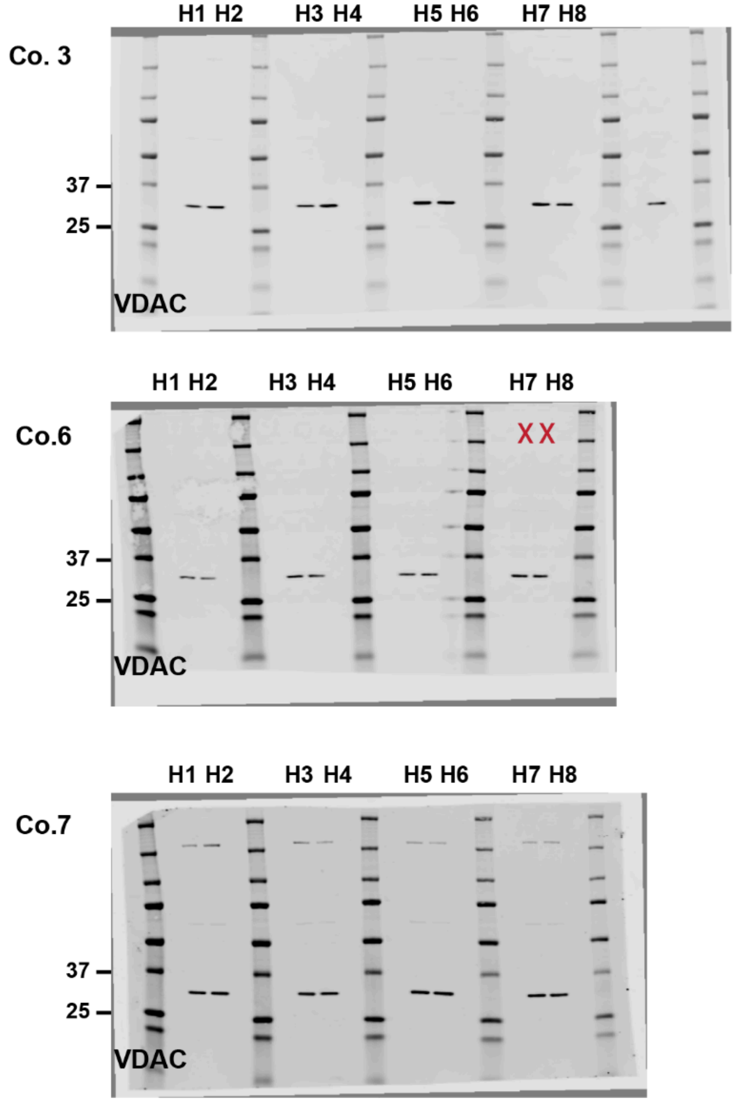


**C**

Cohort 3

| Lane | Ex. ID |
|---|---|
| H1 | G4 -3 |
| H2 | G4 -4 |
| H3 | Td565 |
| H4 | G5-3 |
| H5 | B008 |
| H6 | B005 |
| H7 | B007 |
| H8 | B006 |

Cohort 6

| Lane | Ex. ID |
|---|---|
| H1 | B036 |
| H2 | B026 |
| H3 | B035 |
| H4 | B038 |
| H5 | SS5 |
| H6 | SS4 |
| H7 | Excluded |
| H8 | Excluded |

Cohort 7

| Lane | Ex. ID |
|---|---|
| H1 | B070 |
| H2 | B071 |
| H3 | B074 |
| H4 | B072 |
| H5 | B080 |
| H6 | B081 |
| H7 | B085 |
| H8 | B086 |

**Supplemental Figure 9.** *Total OXPHOS integrated density over VDAC across cohorts*

A. Quantification shows the integrated density of ETC complex I-V subunits relative to loading control VDAC from mitochondrial fractions overexpressing GFP or MCU across cohorts.

B. Uncropped immunoblot stained with antibody against loading control VDAC in mitochondrial fractions isolated from hippocampi overexpressing GFP or MCU. Red X denotes samples excluded from the study.

C. Key provided for each experimental ID across cohorts.

A

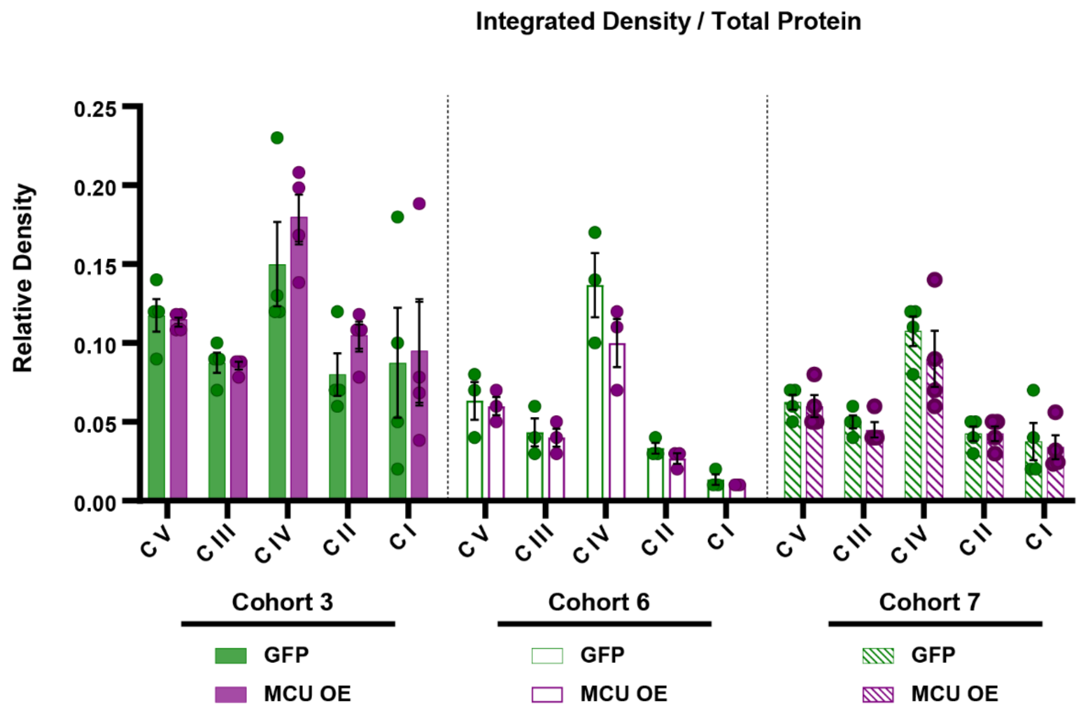


B

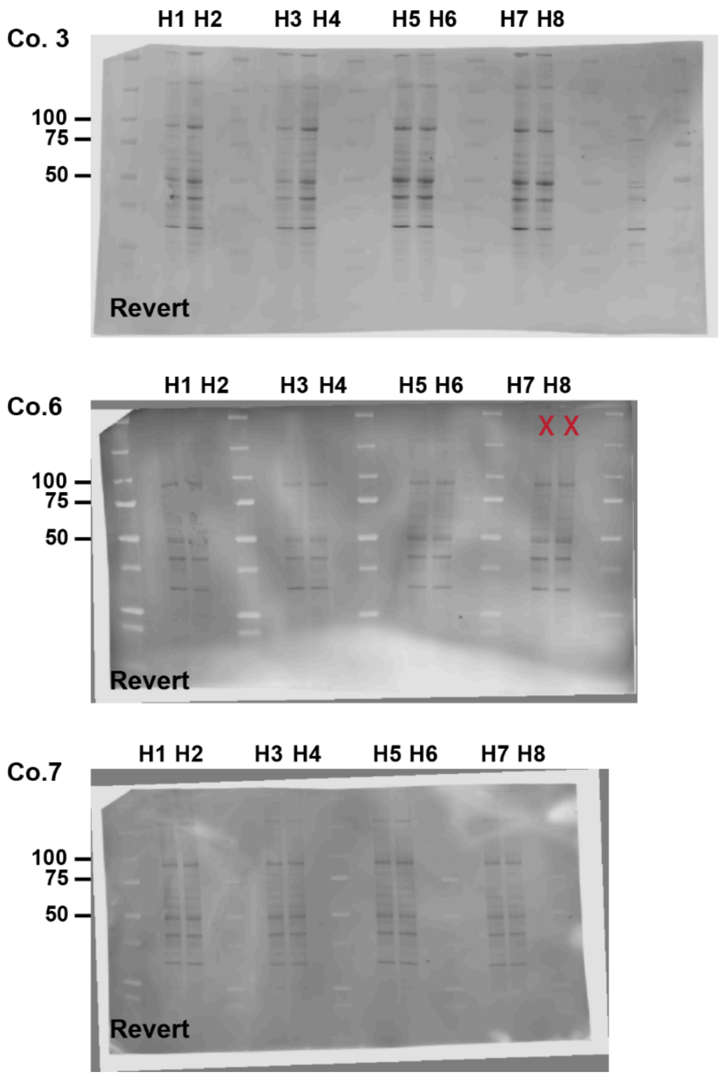


C

Cohort 3

| Lane | Ex. ID |
|---|---|
| H1 | G4 -3 |
| H2 | G4 -4 |
| H3 | Td565 |
| H4 | G5-3 |
| H5 | B008 |
| H6 | B005 |
| H7 | B007 |
| H8 | B006 |

Cohort 6

| Lane | Ex. ID |
|---|---|
| H1 | B036 |
| H2 | B026 |
| H3 | B035 |
| H4 | B038 |
| H5 | SS5 |
| H6 | SS4 |
| H7 | Excluded |
| H8 | Excluded |

Cohort 7

| Lane | Ex. ID |
|---|---|
| H1 | B070 |
| H2 | B071 |
| H3 | B074 |
| H4 | B072 |
| H5 | B080 |
| H6 | B081 |
| H7 | B085 |
| H8 | B086 |

**Supplemental Figure 10.** *Total OXPHOS integrated density over total protein across cohorts*

A. Quantification shows the integrated density of ETC complex I-V subunits relative to total protein from mitochondrial fractions overexpressing GFP or MCU across cohorts.

B. uncropped immunoblot stained for total protein (Revert$^{TM}$) in mitochondrial fractions isolated from hippocampi overexpressing GFP or MCU. Red X denotes samples excluded from the study.

C. Key provided for each experimental ID across cohorts.

A.

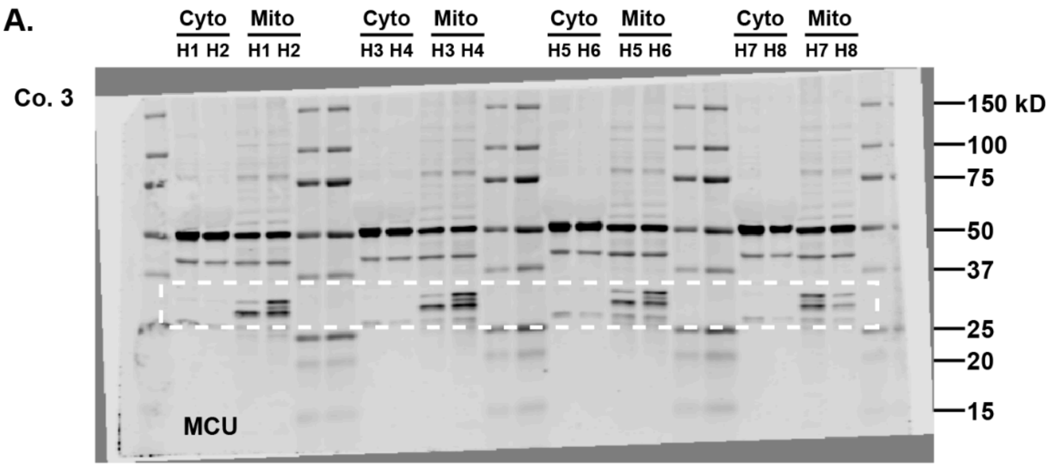


B.

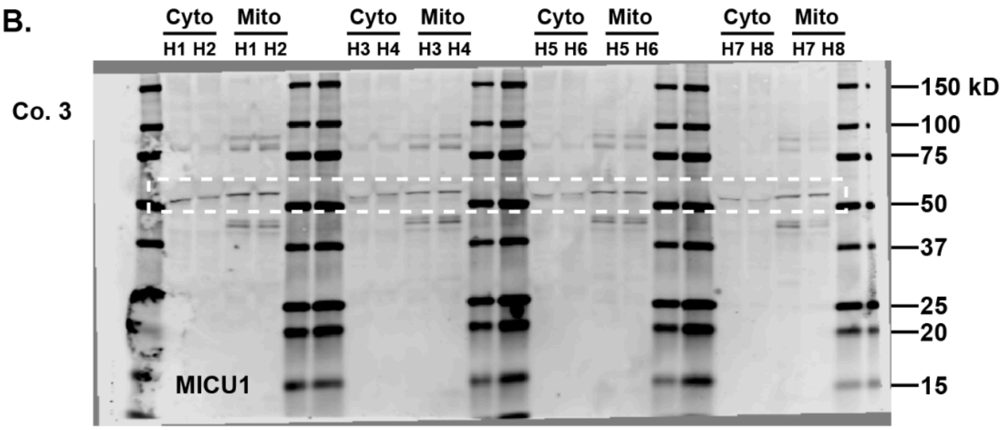


C.

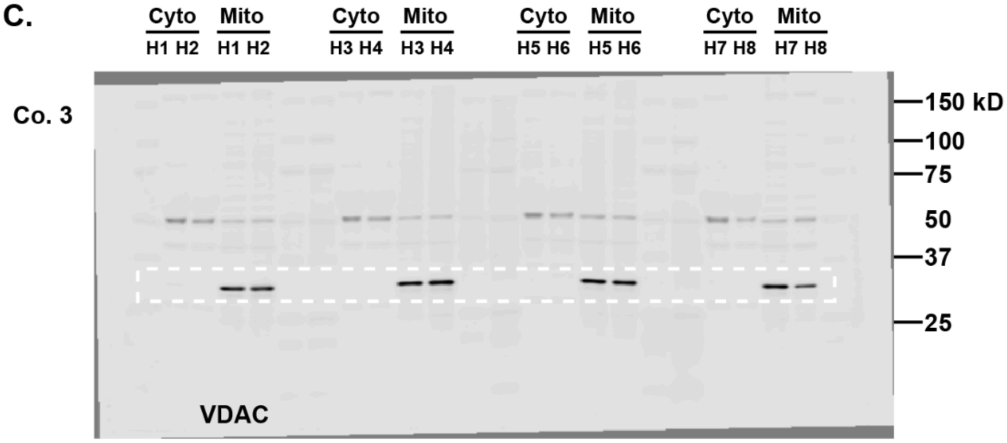


D.

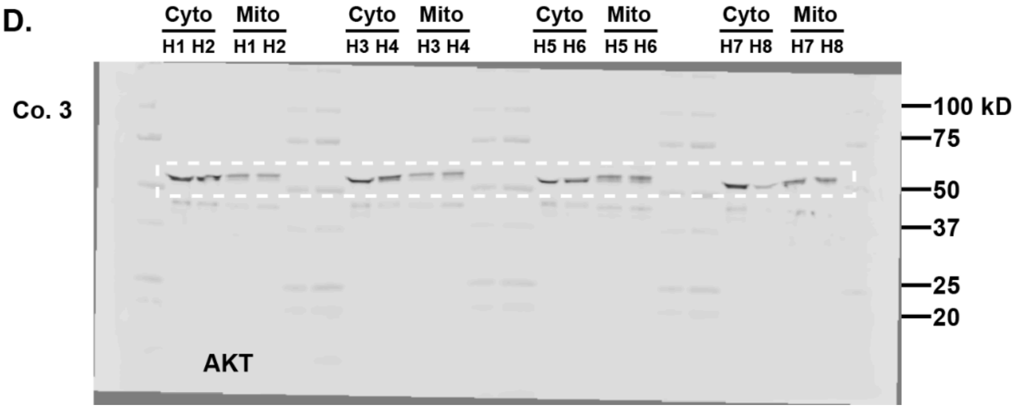


E.

Cohort 3

| Lane | Ex. ID |
|---|---|
| H1 | G4 -3 |
| H2 | G4 -4 |
| H3 | Td565 |
| H4 | G5-3 |
| H5 | B008 |
| H6 | B005 |
| H7 | B007 |
| H8 | B006 |

**Supplemental Figure 11.** *Whole Immunoblots for the representative Western blot image in Figure 1.*

A. Uncropped immunoblot stained with antibodies against MCU (white box) in mitochondrial (mito) and cytosolic (cyto) fractions isolated from hippocampi overexpressing GFP or MCU from cohort 3. Densitometry was used to quantify the total band signal for MCU, which reflected the sum density of multiple bands, and was normalized to the corresponding VDAC loading control to obtain relative MCU expression.

B. Uncropped immunoblot stained with antibodies against MICU1 in mitochondrial and cytosolic fractions isolated from hippocampi overexpressing GFP or MCU.

C. Uncropped immunoblot stained with antibodies against VDAC in mitochondrial and cytosolic fractions isolated from hippocampi overexpressing GFP or MCU. VDAC was used as a loading control to quantify MCU and MICU1 overexpression in mitochondrial fractions.

A) Uncropped immunoblot stained with antibodies against AKT in mitochondrial and cytosolic fractions isolated from hippocampi overexpressing GFP or MCU. AKT was used as a loading control for cytosolic fractions.

B) Key provided for each experimental ID across cohorts.

A.

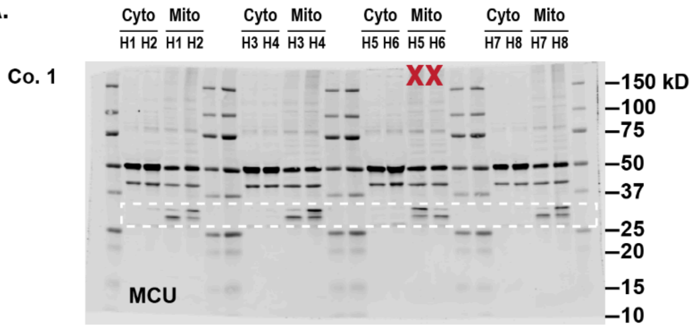


B.

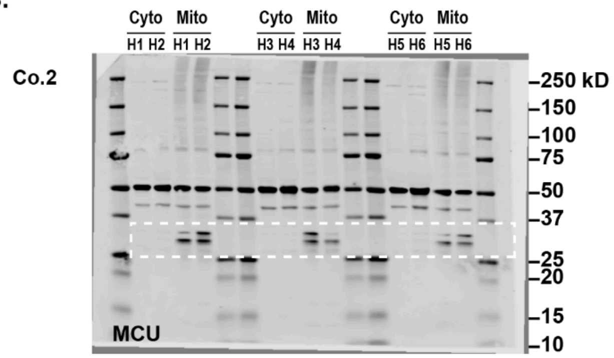


C.

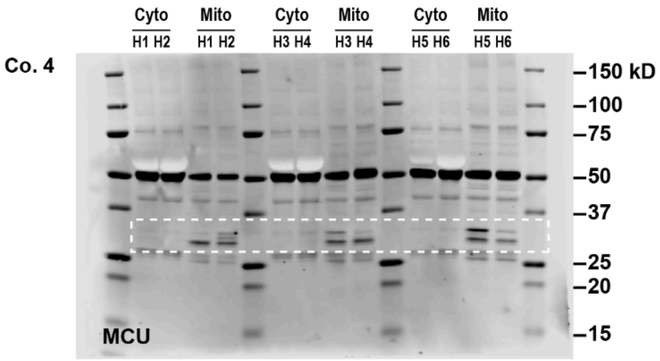


D.

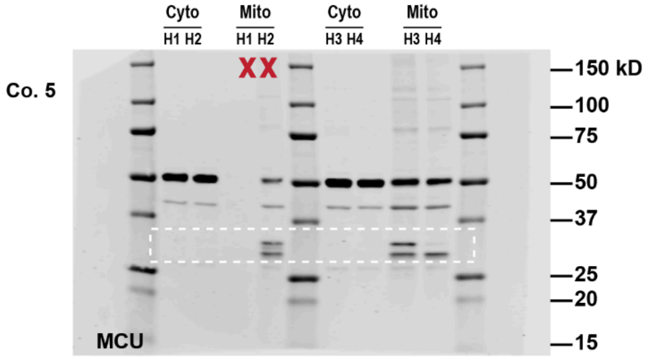


E.

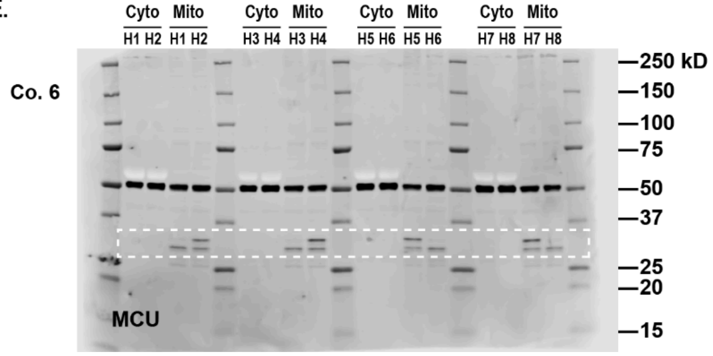


F.

Cohort 1

| Lane | Ex. ID |
|---|---|
| H1 | G823 |
| H2 | Td490 |
| H3 | Td516 |
| H4 | Td515 |
| H5 | Excluded |
| H6 | Excluded |
| H7 | M226 |
| H8 | M227 |

G.

Cohort 2

| Lane | Ex. ID |
|---|---|
| H1 | M229 |
| H2 | M228 |
| H3 | M237 |
| H4 | M238 |
| H5 | M239 |
| H6 | M240 |

H.

Cohort 4

| Lane | Ex. ID |
|---|---|
| H1 | G3-2 |
| H2 | G3-1 |
| H3 | G4-2 |
| H4 | G4-1 |
| H5 | G12-1 |
| H6 | G12-2 |

I.

Cohort 5

| Lane | Ex. ID |
|---|---|
| H1 | Excluded |
| H2 | Excluded |
| H3 | B017 |
| H4 | B018 |

J.

Cohort 6

| Lane | Ex. ID |
|---|---|
| H1 | B036 |
| H2 | B026 |
| H3 | B035 |
| H4 | B038 |
| H5 | SS5 |
| H6 | SS4 |
| H7 | SS7 |
| H8 | SS6 |

**Supplemental Figure 12.** *Uncropped immunoblots for quantification of MCU expression levels.*

A-E) Uncropped immunoblot stained with antibodies against MCU (white box) in mitochondrial (mito) and cytosolic (cyto) fractions isolated from hippocampi overexpressing GFP or MCU. Densitometry was used to quantify the total band signal for MCU, which reflected the sum density of multiple bands, and was normalized to the corresponding VDAC loading control to obtain relative MCU expression. Each gel represents an experimental cohort. Red “X” notes a sample excluded from the analysis.

F-J) Experimental IDs for each cohort associated with the membranes on the left.

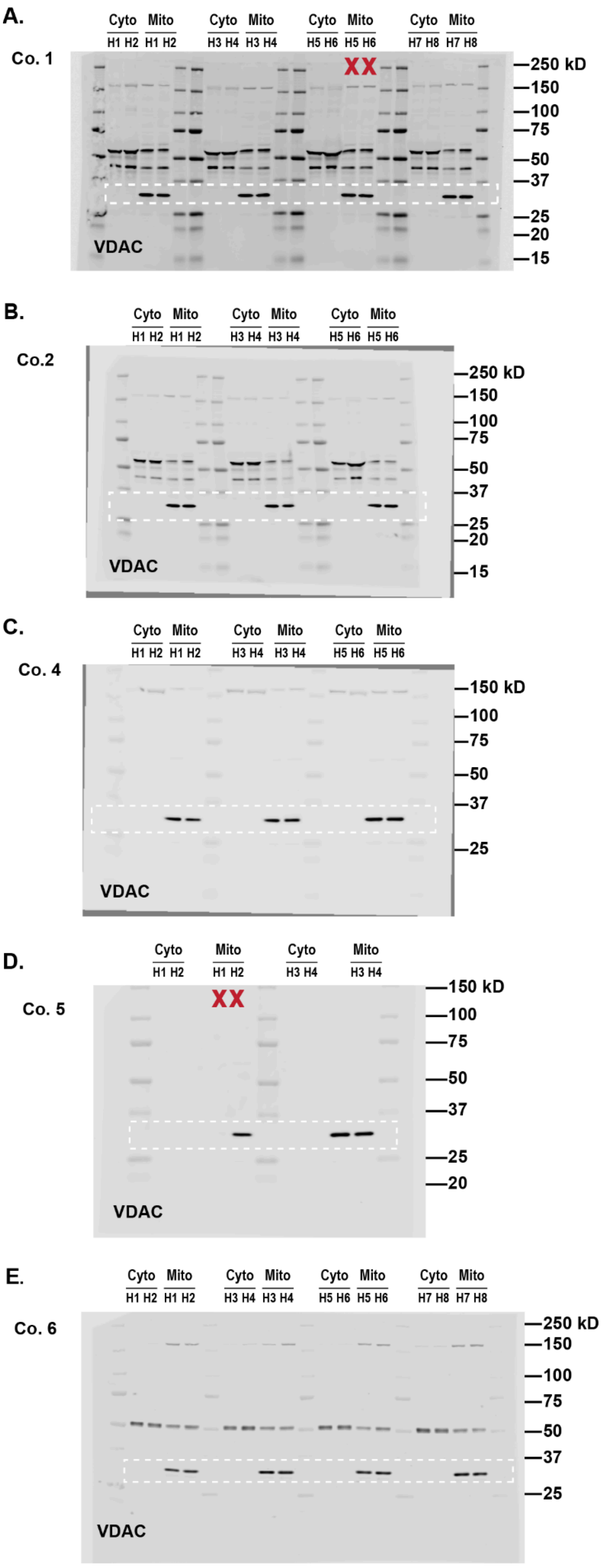


**F.**

Cohort 1

| Lane | Ex. ID |
|---|---|
| H1 | G823 |
| H2 | Td490 |
| H3 | Td516 |
| H4 | Td515 |
| H5 | Excluded |
| H6 | Excluded |
| H7 | M226 |
| H8 | M227 |

**G.**

Cohort 2

| Lane | Ex. ID |
|---|---|
| H1 | M229 |
| H2 | M228 |
| H3 | M237 |
| H4 | M238 |
| H5 | M239 |
| H6 | M240 |

**H.**

Cohort 4

| Lane | Ex. ID |
|---|---|
| H1 | G3-2 |
| H2 | G3-1 |
| H3 | G4-2 |
| H4 | G4-1 |
| H5 | G12-1 |
| H6 | G12-2 |

**I.**

Cohort 5

| Lane | Ex. ID |
|---|---|
| H1 | Excluded |
| H2 | Excluded |
| H3 | B017 |
| H4 | B018 |

**J.**

Cohort 6

| Lane | Ex. ID |
|---|---|
| H1 | B036 |
| H2 | B026 |
| H3 | B035 |
| H4 | B038 |
| H5 | SS5 |
| H6 | SS4 |
| H7 | SS7 |
| H8 | SS6 |

**Supplemental Figure 13.** *Uncropped immunoblot for loading control VDAC*

A-E) Uncropped immunoblot stained with antibodies against loading control VDAC (white box) in mitochondrial (mito) and cytosolic (cyto) fractions isolated from hippocampi overexpressing GFP or MCU. Each gel represents an experimental cohort. Red “X” notes a sample excluded from the analysis.

F-J) Experimental IDs for each cohort associated with the membranes on the left.

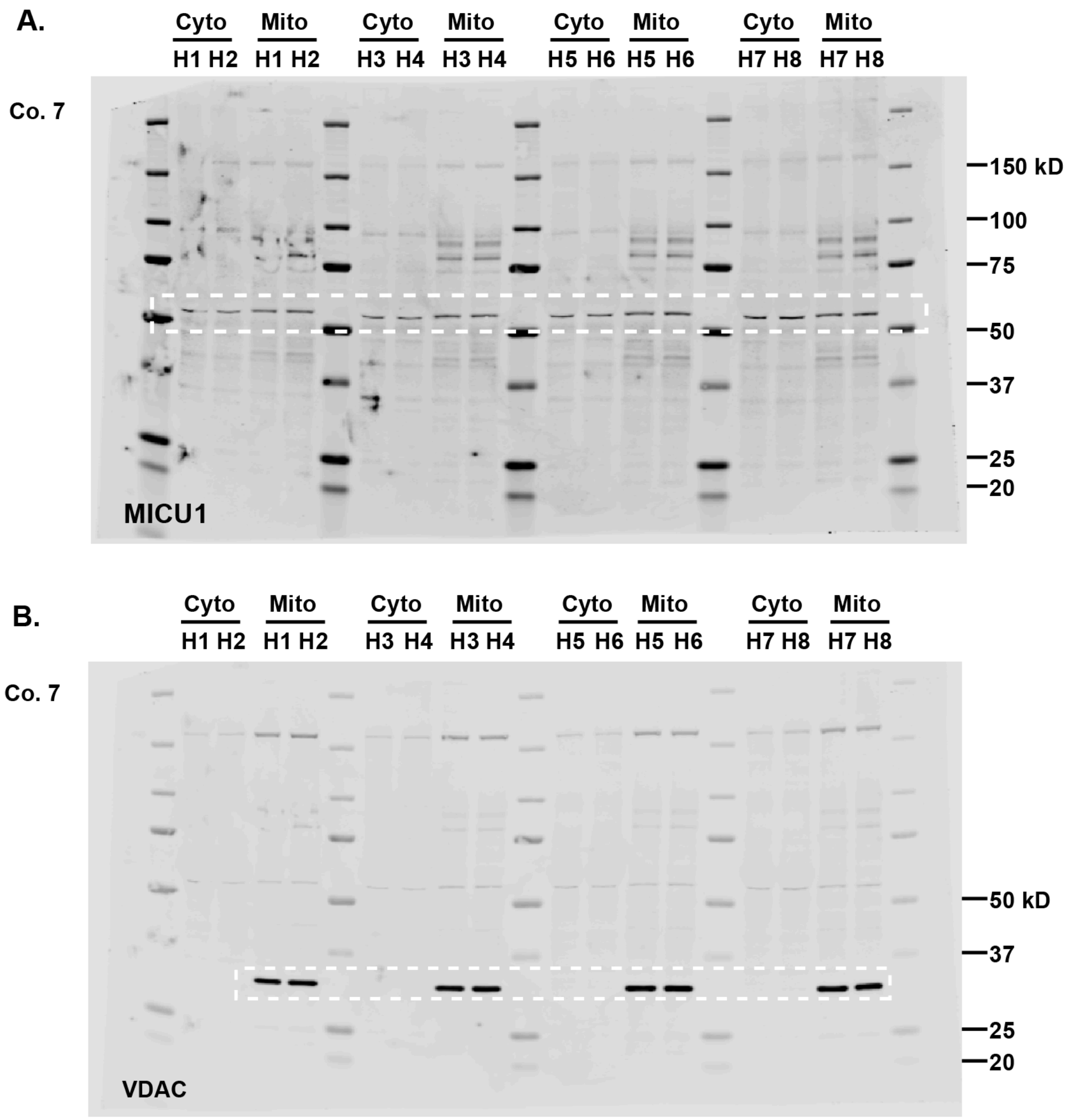


C.

Cohort 7

| Lane | Ex. ID |
|---|---|
| H1 | B070 |
| H2 | B071 |
| H3 | B074 |
| H4 | B072 |
| H5 | B080 |
| H6 | B081 |
| H7 | B085 |
| H8 | B086 |

**Supplemental Figure 14.** *Uncropped immunoblot for MICU1 and loading control VDAC*

A. Uncropped immunoblot stained with antibody against MICU1 (white box) in mitochondrial (mito) and cytosolic (cyto) fractions isolated from hippocampi overexpressing GFP or MCU.

B. Uncropped immunoblot stained with antibody against loading control VDAC (white box) in mitochondrial (mito) and cytosolic (cyto) fractions isolated from hippocampi overexpressing GFP or MCU.

C. Experimental IDs for each cohort associated with the membranes on the left.
