## Supplemental Table 1 for "Elevating levels of neuronal MCU in the hippocampus enhances mitochondrial calcium uptake and respiratory efficiency proportional to demand"

| Cohort | Lane I | Strain | Sex | Group | Mouse ID | Respirati | Calcium | Swelling | MCU OE WI | MICU1 W | Total Oxphos W | TEM | Notes |
|---|---|---|---|---|---|---|---|---|---|---|---|---|---|
| Cohort 1 | H1 | Am2-eGFP | Male | GFP | G823 | O2k | CRC | mPTP | MCU OE | | | | |
| Cohort 1 | H2 | TDtom | Male | MCU | Td490 | O2k | CRC | mPTP | MCU OE | | | | |
| Cohort 1 | H3 | TDtom | Female | GFP | Td516 | O2k | CRC | | MCU OE | | | | |
| Cohort 1 | H4 | TDtom | Female | MCU | Td515 | O2k | CRC | | MCU OE | | | | |
| Cohort 1 | H5 | TDtom | Male | MCU | Td513 | | CRC | | MCU OE | | | | |
| Cohort 1 | H6 | C57bl6 /N | Female | WT | M245 | | CRC | | MCU OE | | | | Cohort 1 H5, H6 - Pair was excluded from CRC experiments and MCU OE quantification because unblinding revealed MCU OE and WT pairing, not a GFP-expressing CTL |
| Cohort 1 | H7 | C57bl6 /N | Female | GFP | M226 | O2k | CRC | | MCU OE | | | | |
| Cohort 1 | H8 | C57bl6 /N | Female | MCU | M227 | O2k | CRC | | MCU OE | | | | |
| Cohort 2 | H1 | C57bl6 /N | Female | GFP | M229 | O2k | CRC | | MCU OE | | | | Cohort 2 H1, H2 - Pair excluded from O2k experiments because state 1 negative OCR, likely indicating poor instrument calibration |
| Cohort 2 | H2 | C57bl6 /N | Female | MCU | M228 | O2k | CRC | | MCU OE | | | | |
| Cohort 2 | H3 | C57bl6 /N | Female | MCU | M237 | O2k | CRC | | MCU OE | | | | Cohort 2 H3, H4 - Pair excluded from CRC experiments due to failed CRC assay and not enough mitochondria to repeat the experiment |
| Cohort 2 | H4 | C57bl6 /N | Female | GFP | M238 | O2k | CRC | | MCU OE | | | | |
| Cohort 2 | H5 | C57bl6 /N | Male | GFP | M239 | O2k | CRC | mPTP | MCU OE | | | | Cohort 2 H5, H6 - Pair excluded from CRC quantification because calcium uptake rates were ≥ 2 SD from sample population |
| Cohort 2 | H6 | C57bl6 /N | Male | MCU | M240 | O2k | CRC | mPTP | MCU OE | | | | |
| Cohort 3 | H1 | C57bl6 /J | Female | GFP | G4 -3 | O2k | CRC | | MCU OE | MICU1 | Total Oxphos | | |
| Cohort 3 | H2 | C57bl6 /J | Female | MCU | G4 -4 | O2k | CRC | | MCU OE | MICU1 | Total Oxphos | | |
| Cohort 3 | H3 | TDtom | Female | GFP | Td565 | O2k | CRC | | MCU OE | MICU1 | Total Oxphos | | |
| Cohort 3 | H4 | TDtom | Female | MCU | G5-3 | O2k | CRC | | MCU OE | MICU1 | Total Oxphos | | |
| Cohort 3 | H5 | C57bl6 /J | Female | GFP | B008 | | | | MCU OE | MICU1 | Total Oxphos | | |
| Cohort 3 | H6 | C57bl6 /J | Female | MCU | B005 | | | | MCU OE | MICU1 | Total Oxphos | | |
| Cohort 3 | H7 | C57bl6 /J | Female | MCU | B007 | | | | MCU OE | MICU1 | Total Oxphos | | |
| Cohort 3 | H8 | C57bl6 /J | Female | GFP | B006 | | | | MCU OE | MICU1 | Total Oxphos | | |
| Cohort 4 | H1 | C57bl6 /J | Male | GFP | G3-2 | O2k | CRC | | MCU OE | | | | |
| Cohort 4 | H2 | C57bl6 /J | Male | MCU | G3-1 | O2k | CRC | | MCU OE | | | | |
| Cohort 4 | H3 | C57bl6 /J | Female | MCU | G4-2 | O2k | CRC | | MCU OE | | | | |
| Cohort 4 | H4 | C57bl6 /J | Female | GFP | G4-1 | O2k | CRC | | MCU OE | | | | |
| Cohort 4 | H5 | C57bl6 /J | Female | MCU | G12-1 | | | | MCU OE | | | | |
| Cohort 4 | H6 | C57bl6 /J | Female | GFP | G12-2 | | | | MCU OE | | | | |
| Cohort 5 | H1 | C57bl6 /J | Female | GFP | B015 | | | mPTP | MCU OE | | | TEM | Cohort 5 H1, H2 - Pair excluded from MCU OE quantification due to low recovery and no response to mPTP assay |
| Cohort 5 | H2 | C57bl6 /J | Female | MCU | B016 | | | mPTP | MCU OE | | | TEM | |
| Cohort 5 | H3 | C57bl6 /J | Female | MCU | B017 | | | mPTP | MCU OE | | | TEM | |
| Cohort 5 | H4 | C57bl6 /J | Female | GFP | B018 | | | mPTP | MCU OE | | | TEM | |

| Cohort | Lane I | Strain | Sex | Group | Mouse ID | Respirati | Calcium | Swelling | MCU OE WI | MICU1 W | Total Oxphos W | TEM | Notes |
|---|---|---|---|---|---|---|---|---|---|---|---|---|---|
| Cohort 6 | H1 | C57bl6 /J | Female | GFP | B036 | | | mPTP | MCU OE | | Total Oxphos | TEM | |
| Cohort 6 | H2 | C57bl6 /J | Female | MCU | B026 | | | mPTP | MCU OE | | Total Oxphos | TEM | |
| Cohort 6 | H3 | C57bl6 /J | Female | GFP | B035 | | | mPTP | MCU OE | | Total Oxphos | TEM | |
| Cohort 6 | H4 | C57bl6 /J | Female | MCU | B038 | | | mPTP | MCU OE | | Total Oxphos | TEM | |
| Cohort 6 | H5 | C57bl6 /J | Female | MCU | SS5 | | | mPTP | MCU OE | | Total Oxphos | TEM | |
| Cohort 6 | H6 | C57bl6 /J | Female | GFP | SS4 | | | mPTP | MCU OE | | Total Oxphos | TEM | |
| Cohort 6 | H7 | C57bl6 /J | Female | MCU | SS7 | | | mPTP | MCU OE | | Total Oxphos | TEM | Cohort 6 H7, H8 - Pair excluded from total oxphos quantification due to gel/image artifact at CI and CII band |
| Cohort 6 | H8 | C57bl6 /J | Female | GFP | SS6 | | | mPTP | MCU OE | | Total Oxphos | TEM | |
| Cohort 7 | H1 | C57bl6 /J | Male | MCU | B070 | | | mPTP | | MICU1 | Total Oxphos | TEM | Cohort 7 H1, H2 - Pair excluded from mPTP quantification due to failed mPTP assay, or no increase in extramitochondrial calcium release |
| Cohort 7 | H2 | C57bl6 /J | Male | GFP | B071 | | | mPTP | | MICU1 | Total Oxphos | TEM | |
| Cohort 7 | H3 | C57bl6 /J | Male | GFP | B074 | | | mPTP | | MICU1 | Total Oxphos | TEM | |
| Cohort 7 | H4 | C57bl6 /J | Male | MCU | B072 | | | mPTP | | MICU1 | Total Oxphos | TEM | |
| Cohort 7 | H5 | C57bl6 /J | Female | MCU | B080 | | | mPTP | | MICU1 | Total Oxphos | TEM | |
| Cohort 7 | H6 | C57bl6 /J | Female | GFP | B081 | | | mPTP | | MICU1 | Total Oxphos | TEM | |
| Cohort 7 | H7 | C57bl6 /J | Female | MCU | B085 | | | mPTP | | MICU1 | Total Oxphos | TEM | |
| Cohort 7 | H8 | C57bl6 /J | Female | GFP | B086 | | | mPTP | | MICU1 | Total Oxphos | TEM | |
